# Cold Exposure Protects from Neuroinflammation Through Immunologic Reprogramming

**DOI:** 10.1101/2020.08.28.269563

**Authors:** Martina Spiljar, Karin Steinbach, Dorothée Rigo, Nicolas Suárez-Zamorano, Ingrid Wagner, Noushin Hadadi, Ilena Vincenti, Nicolas Page, Bogna Klimek, Mary-Aude Rochat, Mario Kreutzfeldt, Claire Chevalier, Ozren Stojanović, Matthias Mack, Dilay Cansever, Melanie Greter, Doron Merkler, Mirko Trajkovski

**Author notes:** Correspondence and equal contribution.

## Abstract

Autoimmunity is energetically costly, but the impact of a metabolically active state on immunity and immune-mediated diseases is unclear. Ly6C^hi^ monocytes are key effectors in CNS autoimmunity with elusive role in priming naïve autoreactive T cells. Here we provide unbiased analysis of the immune changes in various compartments during cold exposure, and show that this energetically costly stimulus markedly ameliorates active experimental autoimmune encephalomyelitis (EAE). Cold exposure decreases MHCII on monocytes at steady-state and in various inflammatory mouse models, and suppresses T cell priming and pathogenicity through the modulation of monocytes. Genetic, or antibody-mediated monocyte depletion, or adoptive transfer of Th1- or Th17-polarized cells for EAE abolish the cold-induced effects on T cells or EAE, respectively. These findings provide a mechanistic link between environmental temperature and neuroinflammation, and suggest competition between cold-induced metabolic adaptations and autoimmunity as energetic trade-off beneficial for the immune-mediated diseases.

## INTRODUCTION

Maintaining immunity requires substantial metabolic resources (Buck et al., 2017; Demas and Nelson, 2012; Ganeshan et al., 2019; Hotamisligil, 2017; McDade, 2005; O’Neill et al., 2016) for steady-state systemic immune surveillance and during inflammation, leading to the generation of millions of immune cells daily. The life-history theory (Stearns, 1992) proposes that prioritization of resources between biological programs would depend on the environment. In hostile environments, resources are shifted away from growth and reproduction programs into maintenance programs (Okin and Medzhitov, 2012; Stearns, 1992; Wang et al., 2019). In addition to immunity, these maintenance traits include adaptations to energy scarcity, but also adaptations to changes in the environmental temperature. Exposure to cold provokes a high cost–high gain physiologic response aimed at reducing heat dissipation (e.g. by vasoconstriction) and increasing heat production, mainly by activating the brown adipose tissue (BAT) and browning of the subcutaneous adipose tissue (SAT). BAT catabolizes energy to enable generation of heat, a function conferred by the uncoupling protein 1 (UCP1). The cold-induced BAT activation is triggered by sympathetic innervation (Cannon and Nedergaard, 2004; Stojanovic et al., 2018). Norepinephrine released from the nerve endings stimulates the beta 3 adrenergic receptor, and agonists of this receptor are frequently used to mimic cold exposure and experimentally induce thermogenesis.

In autoimmunity, the organism develops an energetically costly pro-inflammatory immune response (Okin and Medzhitov, 2012). Such high energy demanding processes may compete with each other, and prioritization of one task over the other may be a result of an energetic trade-off. This concept can be of particular interest for autoimmunity, in which introducing an additional energy-costly program may result in milder immune response and disease outcome. In contrast to the well-described immune status during obesity (Grant and Dixit, 2015; Hotamisligil, 2017; Kohlgruber et al., 2016; Li et al., 2020; Man et al., 2017), the repercussions of a metabolically active phenotype on the autoimmunity, and the potential systemic effects on the immune-mediated diseases are unknown.

Experimental Autoimmune Encephalomyelitis (EAE) is an animal model for multiple sclerosis (MS), the most frequent autoimmune demyelinating disease of the central nervous system (CNS). Similar to MS, EAE is exacerbated by high caloric diet and obesity (Hasan et al., 2016; Piccio et al., 2008; Timmermans et al., 2014; Winer et al., 2009), notably characterized by a positive energy balance. CD4^+^ T cells and monocytes are important cellular players in the pathogenesis of EAE. Among CD4^+^ T cells, distinct encephalitogenic subsets are essentially involved in disease precipitation and are characterized by certain cytokine signatures, including interleukin17 (IL-17), Granulocyte-Macrophage Colony Stimulating Factor (GM-CSF) and Interferon gamma (IFNg). Monocytes and monocyte-derived cells are crucial for the resulting tissue destruction of the central nervous system (CNS), particularly during the effector phase of EAE (Ajami et al., 2011; Fife et al., 2000; Izikson et al., 2000; King et al., 2009; Mildner et al., 2009; Serbina and Pamer, 2006). Thereby, these monocytes and monocyte-derived cells can express pro-inflammatory INOS (m1-like) or anti-inflammatory ARG1 (m2-like) and in addition can exert antigen presenting function via major histocompatibility complex class II (MHCII) upregulation. The role of monocytes in priming of autoreactive T cells in CNS autoimmunity, and whether cold exposure could affect this interaction remain unclear.

Our study provides an unbiased characterization of the immune system in different compartments in healthy conditions and during CNS autoimmunity following exposure to lower environmental temperature. We demonstrate that cold exposure renders monocytes less activated in the bone marrow, the circulation and the CNS. The change in the monocyte phenotype affects priming of pathogenic T cells during neuroinflammation, resulting in reduced T cell cytokine expression and consequently attenuated EAE. Our data suggest competition between cold exposure and autoimmunity leading to a constrained immune response, which could be of therapeutic importance in neuroinflammation, and potentially other autoimmune diseases.

## RESULTS

### Cold Exposure Modulates Myeloid Cells in Bone Marrow at Steady State

To explore the impact of cold exposure on the immune system, we first focused on the bone marrow where immune cells originate from. We performed RNA sequencing of the bone marrow from mice exposed to 10°C for two weeks following an acclimatization period, and compared them to mice kept at room temperature (Figures 1A and S1A). We shortlisted the significantly different genes (p<0.05) for pathway and gene ontology analyses and considered pathways and terms significantly deregulated or enriched when p<0.05. Among the 5088 enriched terms identified by Metacore Gene Ontology Processes, “Myeloid Leukocyte Activation” was among the top five most enriched terms (Figure 1B). Metacore Metabolic Network analysis identified 41 enriched networks (Figure S1B), whereas Metacore Pathway Maps revealed 885 pathways that were changed. Among the top 10 most significantly regulated ones (Figure S1C), four pathways were consistently changed (>80% of genes in same direction) (Figures 1C and S1D). These were Macrophage and DC shift, G-CSF-induced myeloid differentiation, IFN-alpha/beta signaling and Transcription regulation-granulocyte development, which mainly contain genes important for myeloid cell activation and differentiation.

**Figure 1.**
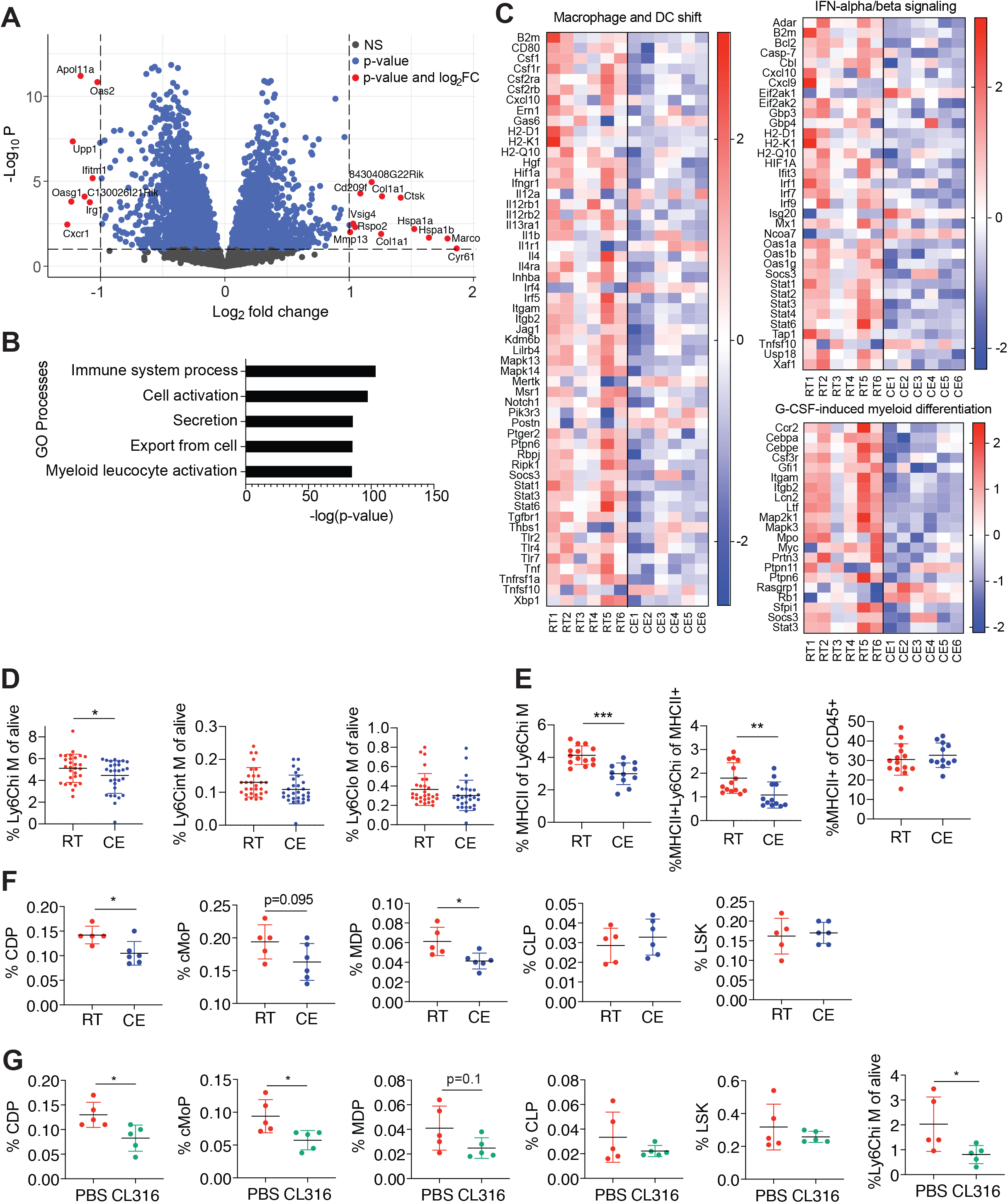
Cold Exposure Impacts Monocytes in the Bone Marrow. (A) Volcano plot showing the up- and downregulated transcripts by RNA sequencing in the bone marrow of cold exposed mice for 2 weeks at 10°C compared to room temperature counterparts (n=6 mice per group). (B) Metacore Gene Ontology Processes analysis displaying the top 5 enriched processes of mice as in (A). (C) Shown are 3 of 4 heatmaps that were regulated in the same direction (> 80% of genes in the same direction) from the top 10 differentially regulated Metacore Pathway Maps of mice as in (A). Full pathway names are “Macrophage and dendritic cell phenotype shift in cancer”, “Immune response_IFN-aplha/beta signaling via JAK/STAT” and “G-CSF induced myeloid differentiation”. (D) Flow cytometry analysis of bone marrow cells of mice as in (A). Percentage of Ly6C high (hi), intermediate (int) and low (lo) monocytes of total, single CD45^+^ alive cells. (E) Percentage of MHCII^+^ cells of Ly6C^hi^ monocytes (left), MHCII^+^ Ly6C^hi^ monocytes of total MHCII^+^ cells (middle) and total MHCII^+^ cells of CD45^+^ cells of mice as in (A), as determined by flow cytometry. (F) Bone marrow immune cell progenitors of mice as in (A) were analyzed by flow cytometry and percentage of total alive single CD45+ cells is shown for common dendritic cell progenitors (CDP), cMoP (common monocyte progenitors), monocyte-dendritic cell progenitors (MDP), common lymphoid progenitors (CLP) and Lin^-^ Sca1^+^ c-KIT^+^ cells (LSK), as determined by flow cytometry. (G) Immune cell progenitor flow cytometry analysis of mice that were i.p. injected with beta 3 adrenoreceptor agonist CL316,243 (CL316) or vector (PBS) daily during one week. Percentage of total alive CD45^+^ cells is shown of CDP, cMoP, MDP, CLP, LSK and Ly6C^hi^ monocytes. (B-C) The cutoff on differentially regulated genes considered for the pathway analysis is p<0.05 and pathways are considered deregulated with p<0.05. Shown are –log(p-value). (D-G) Each dot represents one animal. Shown is mean ±SD. Significance was calculated using student’s t test: ^∗^p < 0.05, ^∗∗^p < 0.01, ^∗∗∗^p < 0.001. Pool of five experiments (D), two experiments (E), one representative experiment of three out of five that were similar (F) and one representative of two are shown.

We then examined the impact of cold exposure at cellular and protein level within the bone marrow by flow cytometry. While cold exposure did not affect the number of total bone marrow cells (Figure S1E), there was a decrease in Ly6C^hi^ but not Ly6C^int^ or Ly6C^lo^ monocytes (Figure 1D). Ly6C^hi^ monocytes showed phenotypical changes characterized by decreased MHCII expression (Figure 1E), a marker for antigen presentation and activation. Within the total MHCII^+^ cells, the population of MHCII^+^Ly6C^hi^ monocytes was decreased, while the percentage of total MHCII^+^ cells within the bone marrow remained unchanged (Figure 1E, gating strategy S1F). Differences in the MHCII expression on protein level were consistent with the observations from RNA sequencing, where MHCII genes were within the top downregulated pathways (Figure 1C).

To address whether cold exposure decreases the monocytes by affecting hematopoiesis, we analyzed hematopoietic progenitors in the bone marrow similarly as previously described (Hettinger et al., 2013; Luo et al., 2015; Merad et al., 2013). In three out of five experiments, cold exposure decreased the absolute cell numbers (data not shown) and the percentage of monocyte-dendritic cell progenitors (MDPs), common dendritic cell progenitors (CDP) and tendentially common monocyte progenitors (cMoP), while the common lymphoid progenitors (CLP) and most hematopoietic cells (Lin- Sca1+ c-KIT+ or LSK) remained unchanged (Figure 1F, gating strategy S1F).

Beta 3 adrenergic receptor signaling is one of the main pathways activated by cold exposure (Cannon and Nedergaard, 2004; Stojanovic et al., 2018). The beta 3 adrenoreceptor agonist CL316,243 (CL316) decreased Ly6C^hi^ monocytes, common dendritic cell progenitors and common monocyte progenitors (Figure 1G), suggesting that beta 3 adrenoreceptor stimulation is sufficient to partly mimic the effects of cold on the myeloid bone marrow cells and their progenitors. Together, these results suggest that cold exposure decreases myeloid cell progenitors and monocytes in the bone marrow, and downregulates monocyte MHCII expression.

### Cold Exposure Impacts Monocytes in Blood at Steady State

We next examined the systemic relevance of the cold exposure-mediated changes observed in the bone marrow by FACS sorting CD115^+^ blood monocytes (Figure S2A) and subsequent profiling them via RNA sequencing (Figures S2B and 2A). Among the 58 Gene Ontology Biological Processes that were enriched, the top 15 terms included leukocyte migration, T cell co-stimulation and antigen presentation via MHCII (Figure 2B), all processes important for the immune system in health and disease. Such pathways were similarly found within the top 15 of 653 total Metacore Pathway Maps (no p-value cutoff) (Figure S2C). Metacore Metabolic Networks analysis revealed that 13 metabolic terms were enriched (Figure S2D). Among the variety of general cell membrane restructuring elements (p<0.05, total 12), “MHC class II protein complex” was among the most enriched Cellular Components in Gene Ontology analysis (Figure S2E). To further profile the effect of cold exposure on circulatory immune cells and in particular myeloid cells, we performed high parameter flow cytometry (BD FACSymphony), followed by an unbiased analysis using FlowSOM algorithm (Van Gassen et al., 2015) and visualization via UMAP plots. Overall, we neither observed any major shifts in myeloid cell (Figure 2C), nor in T cell populations (data not shown) or numbers, suggesting that cold exposure induces several gene expression changes on the monocytes without significantly altering the overall immune cell composition. These observations were further supported by flow cytometry, which revealed that cold exposure consistently decreased MHCII expression of Ly6C^hi^ monocytes, without altering their overall number (Figures 2D and 2E, gating strategy Figure S2F). Similarly, as in the bone marrow, beta 3 adrenoreceptor stimulation mimicked the effects of cold exposure and reduced the MHCII expression of Ly6C^hi^ monocytes (Figure 2F). These data indicate that cold exposure induces functional changes on the monocytes in the blood by reducing their MHCII expression, without provoking major changes in the main immune cell populations.

**Figure 2.**
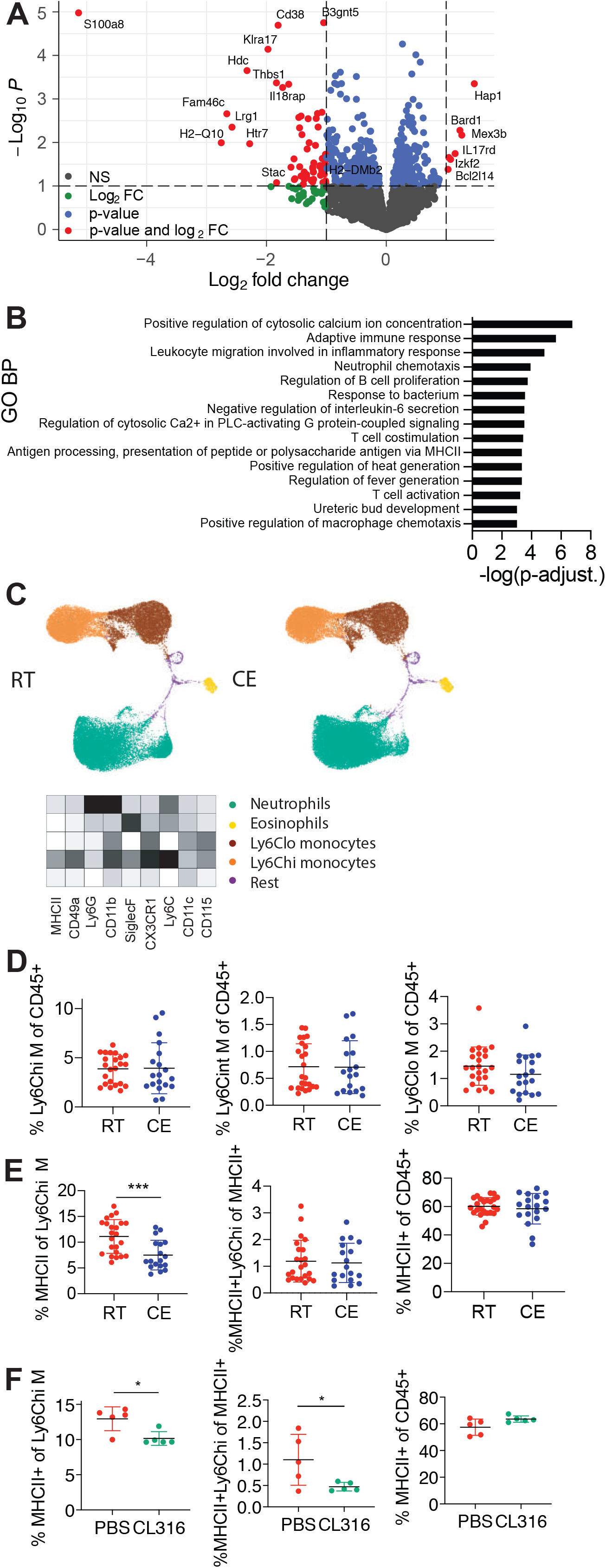
Cold Exposure Modulates Monocytes in the Circulation. (A) Volcano plot showing the up- and downregulated transcripts by RNA sequencing of monocytes that were MACS (anti-PE) and FACS (CD115-PE+) sorted from blood of 2 weeks cold exposed (10°C) mice compared to room temperature mice. (B) 15 most enriched Gene Ontology Biological Processes of monocytes as in (A). Genes were considered differentially regulated with p<0.05. Shown are –log(p-value). (C) Blood immune cells of mice as in (A) visualized using UMAP and clustered using FlowSOM algorithm in R. Ly6C^hi^ monocytes (green), Ly6C^lo^ monocytes (yellow), neutrophils (brown), eosinophils (orange) and others. Heatmap shows median relative expression of all panel markers. (D) Flow cytometry analysis of blood cells from mice as in (A). Percentage of Ly6C high (hi), intermediate (int) and low (lo) monocytes of total, single, CD45^+^ cells. (E) Percentage of blood MHCII^+^ cells of Ly6C^hi^ monocytes (left), MHCII^+^ Ly6C^hi^ monocytes of total MHCII^+^ cells (middle) and total MHCII^+^ cells of total, single CD45^+^ cells of mice as in (A). (F) Blood cell analysis after 1 week of daily i.p. injected beta 3 adrenoreceptor agonist CL316,243 or vector (PBS). Percentage of MHCII^+^ cells of Ly6C^hi^ monocytes (left), MHCII^+^ Ly6C^hi^ monocytes of total MHCII^+^ cells (middle) and total MHCII^+^ cells of CD45^+^ cells. (D-F) Each dot represents one animal. Shown is mean ±SD. Significance was calculated using student’s t test: ^∗^p < 0.05, ^∗∗^p < 0.01, ^∗∗∗^p < 0.001. Pool of three experiments (D, E).

### Cold Decreases Monocyte MHCII Expression in Inflammatory Mouse Models

To study the functional relevance of the observed cold-induced monocyte modulations, we used different immunologic models. First, we amplified circulating monocytes and their MHCII expression using B16-GM-CSF tumor cells (Dranoff et al., 1993; Menezes et al., 2016) that were subcutaneously injected into cold exposed-, or room temperature-kept mice. GM-CSF initiates an inflammatory phenotype in monocytes and depletion of monocyte GM-CSF signaling renders mice resistant to EAE (Croxford et al., 2015). Cold exposure decreased MHCII expression of monocytes without changing the monocyte percentage after implantation of the tumor cells (Figures 3A). To confirm that this effect is monocyte-specific we also analyzed total CD11c^+^ cells, the majority of which are dendritic cells that remained unchanged (Figure 3B). Thus, cold exposure counteracted GM-CSF-induced increase of MHCII on Ly6C^hi^ monocytes.

**Figure 3.**
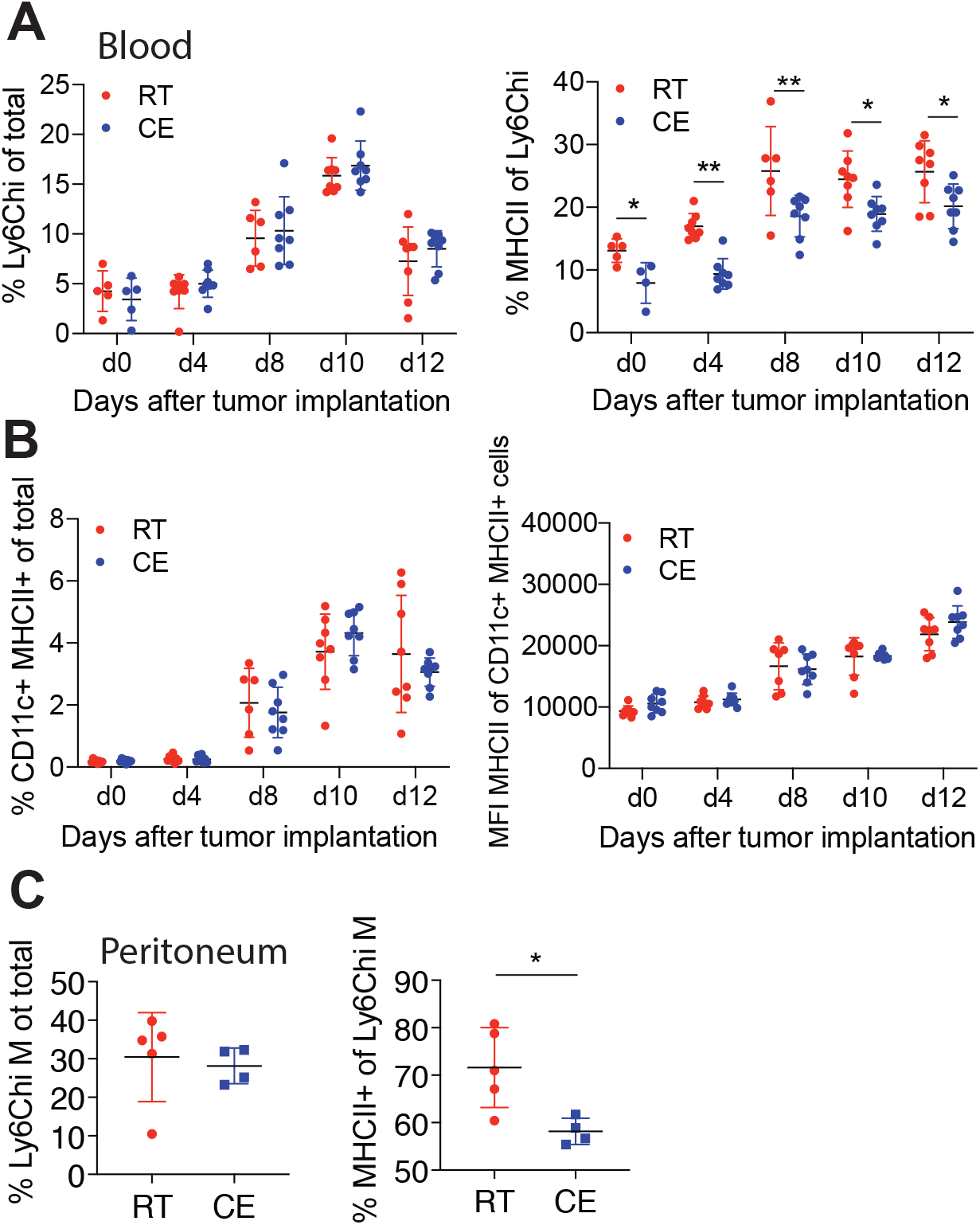
Cold Exposure Decreases Monocyte MHCII in Inflammatory Mouse Models. (A) Blood Ly6C^hi^ monocytes and their MHCII expression were analyzed by flow cytometry on different days after B16-GMCSF s.c. tumor implantation in mice that were cold (10°C) or room temperature exposed for 2 weeks before and during tumor growth. (B) Blood CD11c^+^MHCII^+^ cells and MHCII mean fluorescence intensity (MFI) of MHCII+CD11c^+^ cells were determined by flow cytometry from mice as in (A). (C) Flow cytometry analysis of peritoneal fluid cells 24h after i.p. injection with thioglycollate into 2 weeks cold exposed (10°C) or room temperature mice. (A-C) Each dot represents one animal. Shown is mean ±SD. Significance was calculated using multiple t-test with Holm-Sidak correction (A, B) or student’s t test: (C), ^∗^p < 0.05, ^∗∗^p < 0.01.

Next, we triggered a thioglycolate elicited peritoneal inflammation, in which monocytes are highly attracted into the peritoneum. Cold exposure decreased MHCII expression of recruited Ly6C^hi^ monocytes/monocyte-derived cells within the peritoneum (Figure 3C), without affecting the proportion of several other immune cell populations (Figure S3A). These data show that cold exposure leads to a preferential decrease in the monocyte MHCII expression in various mouse models irrespective of the inflammatory trigger.

Monocytes can acquire antigen presenting functions, including antigen uptake and migration to the lymph nodes (Guilliams et al., 2018; Jakubzick et al., 2017). We therefore investigated if antigen presenting functions of monocytes, as well as dendritic cells (Ganguly et al., 2013), might be changed upon cold exposure. For this purpose, we performed a FITC painting assay, in which FITC is applied epicutaneously of one flank of the mouse and allows measuring migratory immune cell properties into the draining lymph node (dLN) (Allan et al., 2006). We monitored the FITC uptake in the ipsi- and contralateral LN at two different time points, where 12h indicates possible initial differences in lymphatic draining, while 18h is the peak and allows an estimate of the overall changes (Platt et al., 2013). At none of the monitored time points have we observed changes in cell count, FITC uptake or MHCII expression on dendritic cells, monocytes and various dendritic cell subsets (Figure S3B and data not shown), thus excluding that cold exposure affects antigen drainage to the LN. Altogether, these data show that exposure to lower environmental temperature leads to a decrease in the monocyte MHCII expression in various inflammatory mouse models without affecting the lymphatic drainage.

### Cold Exposure Attenuates Experimental Autoimmune Encephalomyelitis (EAE)

Monocytes and monocyte derived cells are important pathogenic players in chronic inflammatory diseases, such as MS. Similarly, monocyte trafficking to the CNS is critical for EAE progression (Ajami et al., 2011; King et al., 2009) and *Ccr2*-deficient mice, which lack monocytes outside the bone marrow (Serbina and Pamer, 2006), show mild (Mildner et al., 2009) or no EAE (Fife et al., 2000; Izikson et al., 2000). We therefore explored whether cold exposure could affect the EAE disease outcome. For this purpose, we exposed mice to 10 °C for 2 weeks with an additional initial acclimatization period of 5 days at 18°C and 5 days at 14°C before immunization, and kept them at 10°C during the whole EAE (Figure 4A). Strikingly, exposing animals to cold led to a pronounced attenuation of the EAE disease severity compared to room temperature kept controls (Figures 4B and S4A). The cold exposed mice also showed delayed onset of the disease, together with a lower maximum and cumulative EAE score, disease incidence and body weight loss (Figures 4C-4G) compared to the RT control mice. Histologic evaluation revealed a reduced area of demyelination in the spinal cord (Figure 4H) and a lower number of lesions (Figure S4B) in spinal cords from cold exposed mice at the peak of EAE. Mac3^+^, CD3^+^ and APP lesions from LFB-Pas per white matter area showed a tendential decrease (Figures S4C-S4G) without reaching statistical significance. As the extent of EAE amelioration slightly differed between experiments, a pool is shown (Figures 4 B-G) of experiments performed under the same conditions except the housing room, which had no consistent impact.

**Figure 4.**
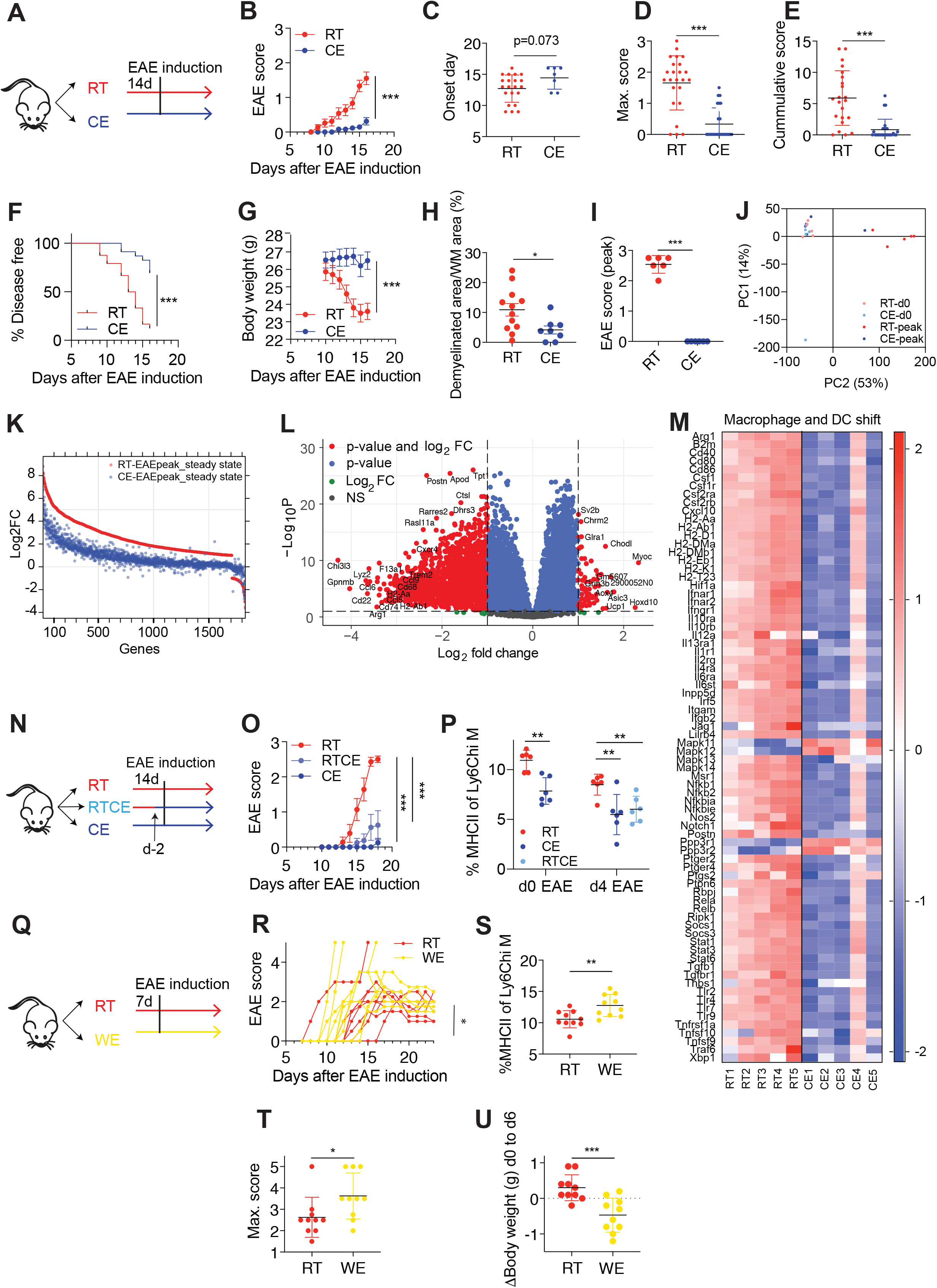
Cold Exposure Attenuates Neuroinflammation. (A-B) Scheme showing the experimental setup for EAE and cold exposure (A). EAE was induced by s.c. immunization with MOG35-55 peptide in complete Freund’s adjuvant (day 0) and pertussis toxin (day 0 and 2). Mice were exposed to cold or room temperature (10°C) for two weeks before and during EAE. Clinical symptoms of EAE were monitored according to a standardized scoring system (B). (C-G) Day of EAE onset, i.e. when first symptoms occur (C), maximum disease score, i.e. the highest score an individual mouse reached during the experiment (D), cumulative disease score, i.e. the sum of all scores each individual mouse reached during the experiment (E), the percentage of disease free mice (F) and body weight (G) of mice as in (A). (H) Quantification of demyelinated area expressed as percentage of white matter (WM) detected by Luxol Fast Blue and Periodic acid–Schiff (LFB-PAS) of spinal cords from mice as in (A). (I-M) Spinal cords of healthy and peak EAE mice that were kept at room temperature or 2 weeks cold exposed were used for RNA sequencing. Scores of the respective mice at sacrifice (I), principal component analysis (J), relativeness analysis of logFC of room temperature healthy vs EAE spinal cords (red) and cold exposure healthy vs EAE spinal cords (blue) with p<0.05, FC>2 (K), volcano plot (gene *Gpx3* was excluded for better visualization) (L) and genes of top1 deregulated pathway “Macrophage and DC phenotype shift in cancer” from Metacore Pathway Maps with p<0.05 (M). (N-O) Scheme showing the experimental setup. Mice were either exposed to cold temperature for two weeks or two days before and continued during EAE and compared to room temperature controls (N). EAE was induced as in (A) and EAE scores are shown (O). (P) Percentage of MHCII expression on Ly6C^hi^ blood monocytes was determined using flow cytometry on day zero and day four after EAE induction. (Q-R) Scheme showing the experimental setup (Q). Mice were exposed to warm temperature of 34°C for one week before and during EAE (as in (A)) and compared to room temperature controls. Each curve represents one individual mouse (R). (S) Percentage of MHCII expression of Ly6C^hi^ blood monocytes from mice as in (Q) was determined using flow cytometry on day 0 of EAE. (T-U) Maximum disease score (T) or delta of body weight gain from day zero to day six of warm exposure (U). (B-S) Data (B-G) represents pool of 3 experiments (n = 8-12 mice per group per experiment). Shown is mean ±SEM, two-way ANOVA (B; G, O, R); student’s t test (C-E, I, S-U); Mantel-Cox (F); multiple t-test with Holm-Sidak correction (P), mean ±SD, ^∗^p < 0.05, ^∗∗^p < 0.01, ^∗∗∗^p < 0.001.

To gain insights into the level of protection by cold exposure on the EAE-induced gene expression alterations in the spinal cord, we performed RNA sequencing on mice kept at room or cold temperature both at steady state, or at EAE peak. Principle component and multidimensional scaling analysis revealed that when the room temperature-kept animals reached peak of the disease after EAE induction (Figure 4I), the immunized cold-exposed mice clustered together with the healthy, steady-state animals (Figures 4J and S4H). Interestingly, all genes that were increased in the room temperature mice with EAE compared to healthy controls were left unchanged, or deregulated to a lower extent in the cold exposed mice under EAE (Figure 4K). These data show that the EAE-provoked spinal cord gene expression alterations are either reduced, or prevented by cold exposure. Gene expression comparison between spinal cords from cold exposed mice vs. room temperature controls during EAE revealed 61 up- and 1579 downregulated genes of 15467 total genes (p<0.05, logFC>1) (Figure 4L). Among the top 984 regulated pathways identified by the Metacore Pathway Maps, “Macrophage and Dendritic cell phenotype shift…”, was the most significantly changed one. This pathway includes MHCII genes and other activation markers (Figures 4M and S4I), further indicating that the myeloid cell and antigen presentation are the most prominent changes induced by cold also during EAE. The pathway further reveals downregulation of CXCL10, a recently identified marker of a pathogenic monocyte subset (Giladi et al., 2020), which we found similarly decreased in the bone marrow (Figure 1C).

To investigate whether the length of the cold exposure before the EAE induction is important for the EAE onset and disease progression, we pre-exposed mice to cold two days before immunization, and compared them to the animals that were kept 2 weeks at 10°C as in the standard protocol. Two days cold pre-exposure was sufficient to attenuate EAE, although to a slightly lower extent than the 2 weeks pre-exposed ones (Figures 4N and 4O), and to reduce monocyte MHCII (Figure 4P), suggesting that the length of the cold pre-treatment plays a modest role in the disease outcome.

To address whether warm exposure has an opposing effect on EAE, we exposed mice to 34°C for seven days before immunization, and kept them at warm during EAE (Figure 4Q). Warm exposure led to decreased food intake and body weight (Figure S4M, Figures 4O and U). Although more mice died under warm exposure, the survival rate was not significantly different, and neither was the day of onset (Figures S4N and S4O). However, in contrast to the effects seen during cold, the MHCII expression of Ly6C^hi^ monocytes was increased in the warm-exposed mice, coupled to deteriorated EAE, enhanced body weight loss and increased maximum disease score (Figures 4R-T and S4J, K). Together, these results suggest that the temperature is a critical determinant for EAE severity, and that cold exposure is a potent stimulus that attenuates neuroinflammation.

Long-term cold exposure increases energy expenditure by enhancing the thermogenesis in the brown and the beige fat, enabling homeothermy (Chevalier et al., 2015). However, when additional biologic maintenance programs are active, as for instance in inflammation, we hypothesized that these responses (maintenance of homeothermy and autoimmunity) may be in competition for energy. We therefore analyzed a variety of parameters involved in thermogenesis in cold the exposed groups of mice. There was no difference in the body temperature between the RT-kept, or cold exposed healthy vs. EAE mice (Figure S5A). While under RT, EAE reduced the blood glucose levels, 3 hour fasting glycemia during cold was unchanged between EAE and healthy mice (Figure S5B). To further address the functional properties of the BAT and SAT during cold, we studied the oxygen consumption rates, which were similar between the cold exposed EAE and healthy mice in both SAT and BAT (Figures S5C, D). Consistent with the lack of functional differences, the weights of the BAT and SAT depots of the EAE mice compared to the healthy cold exposed controls were unchanged (Figures S5E). While the expression of *Cidea* and *Pparg* in the BAT was reduced in the cold exposed EAE mice, the rest of the brown fat markers were unaltered, and there was no difference in expression in any of the browning markers in the inguinal SAT (Figures S5F, G). Moreover, we detected no changes in the lipid droplet size distribution between cold exposed healthy or EAE mice, together suggesting no detectable EAE-induced limitation of the BAT and SAT thermogenic response during cold. However, the slightly milder brown adipose tissue marker activity under EAE, but without overall reduction in the thermogenesis and oxygen consumption rates may suggest competition for energetic resources between of the two energy costly programs. Intriguingly, the lack of functional limitation in the thermogenic response to cold, coupled to decreased neuroinflammation indicates prioritization of the homeothermic response over the autoimmunity, resulting in ameliorated EAE severity.

Cold exposure activates BAT thermogenesis primarily through UCP1, which uncouples the mitochondrial respiratory chain activity from the ATP biosynthesis. However, during prolonged acclimatization to cold, the *Ucp1*-KO mice are able to compensate for the lack of UCP1 through be additional UCP1-independent thermogenic mechanisms that can satisfy the thermogenic needs, but at a very high energetic cost (Ikeda et al., 2017). Surprisingly, *Ucp1* deletion rendered mice even more protected from EAE compared to their wildtype littermates (Figure S5I), associated with increased BW of the *Ucp1 -*KO compared to their WT littermates during cold and EAE, consistent with the overall improved health status. These data demonstrate that the UCP1-dependent BAT activity is dispensable for these effects. They also show that cold exposure ameliorates EAE, but without a cost on the overall homeothermic response.

### Cold Exposure Decreases Monocyte and T Cell Pathogenicity during EAE

Monocytes play an essential role in EAE development, progression and remission (Ajami et al., 2011; Fife et al., 2000; Izikson et al., 2000; King et al., 2009; Mildner et al., 2009; Serbina and Pamer, 2006), and they reach the CNS in inflammatory conditions through the blood stream. To examine whether cold exposure affects monocytes in the circulation during EAE, we performed RNA sequencing on blood monocytes after FACS sorting (Figure S5A, S6A, B) at the EAE onset. Fold change analysis (p<0.05, logFC>1) revealed 660 up- and 86 downregulated genes of 10794 total genes (Figures 5A) in cold exposed compared to room temperature mice. Metacore Metabolic Network analysis (p<0.05) showed 23 enriched networks (Figure 5B). Among the 3366 enriched terms identified using Metacore GO, the top 10 terms were mainly metabolic changes (Figure 5C). Interestingly, Gene Ontology Cellular Components identified “MHC class II protein complex” among the top 5 of 10 total enriched terms (Figure 5B middle). Within the Reactome pathway analysis, pathways important for antigen presentation were within the 4 identified hits (Figure 5B bottom).

**Figure 5.**
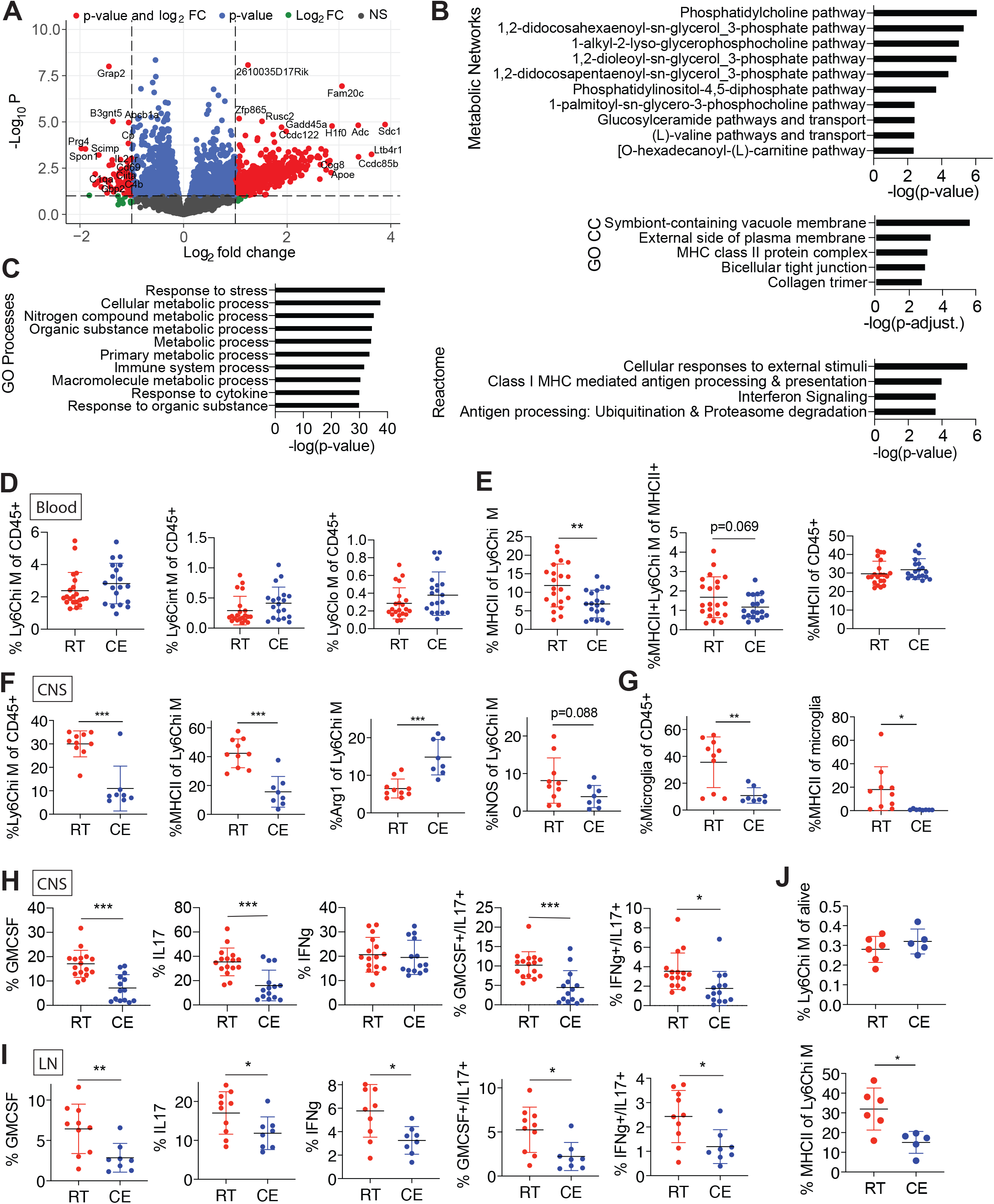
Cold Exposure Reduces Monocyte and T Cell Pathogenicity during EAE. (A) Volcano plot identifying the up- and downregulated transcripts after RNA sequencing of monocytes that were MACS (anti-PE) and FACS (CD115-PE^+^) sorted from blood of mice at onset of EAE (as in Figure 4A) and exposed to cold (10°C) or room temperature for 2 weeks before and continued during EAE. (B-C) Top deregulated and enriched Metacore Metabolic Networks (B top), Gene Ontology cellular components (B middle), Reactome pathways (B bottom) and Gene Ontology Processes (C). Genes were considered when p<0.05 and for Reactome pathways in addition FC>2. Data of sequencing as in (A). (D-E) Flow cytometry analysis of blood cells of mice as in (A) at EAE onset. Percentage of Ly6C high (left), intermediate (middle) and low (right) monocytes of total, single, CD45^+^ alive cells (D). Percentage of MHCII^+^ cells of Ly6C^hi^ monocytes (left), MHCII^+^ Ly6C^+^ monocytes of total MHCII^+^ cells (middle) and total MHCII^+^ cells of CD45^+^ cells (E). Pool of 3 experiments. (F-H) Flow cytometry analysis of CNS cells from mice as in (A) at EAE onset. Percentage of Ly6C^hi^ monocytes/monocyte-derived cells of total, single, CD45^+^ alive cells (F, left). Percentage of MHCII, arginase 1 (Arg1) and iNOS expression of Ly6C^hi^ monocytes/monocyte-derived cells (F, right). Percentage of microglia of total, single, CD45^+^ alive cells and percentage of MHCII of microglia (G). Percentage of corresponding cytokine expression as indicated in CD4^+^ T cells (H). (I) Flow cytometry analysis of dLN cells from mice as in (A) at EAE onset. Percentage of cytokine expression in CD4^+^ T cells. Observed in 2 out of 3 experiments. (J) Flow cytometry analysis of dLN cells from mice as in (A) on day two after EAE induction. Percentage of Ly6C^hi^ monocytes of total, single, CD45^+^ alive cells (upper panel). Percentage of MHCII on Ly6C^hi^ monocytes (lower panel). (D-J) Shown is mean ±SD, significance was calculated using student’s t test, * p<0.05, ** p<0.01, ***p<0.001. Shown is 1 representative example of 3 (F), a pool of 2 experiments (H) or one representative experiment of two out of three that were similar (I).

The changes in MHCII antigen presentation machinery in blood monocytes at EAE onset between the cold exposed and the room temperature mice were confirmed by flow cytometry (Figure S6C). While the percentage of Ly6C^hi^/^int^/^lo^ monocytes remained the same (Figure 5D), cold exposure led to decreased MHCII of Ly6C^hi^ monocytes, coupled to a partial lowering (p=0.069) in the percentage of MHCII^+^Ly6C^hi^ M within the total MHCII^+^ cells but no change in the percentage of total MHCII expression of CD45^+^ cells (Figure 5E). Blood and lymph node immune cells at different days of EAE were further investigated using high dimensional flow cytometry and unbiased computed analysis and detected no alterations thus excluding changes in the myeloid cell populations (Figures S6D and S6E). These analyses demonstrate that cold exposure reduced MHCII expression of circulating Ly6C^hi^ monocytes at EAE onset without shifting myeloid cell populations.

Circulating Ly6C^hi^ monocytes contribute to EAE pathology by migration to the inflamed CNS (Ajami et al., 2011; Fife et al., 2000; Izikson et al., 2000; King et al., 2009; Mildner et al., 2009; Serbina and Pamer, 2006). We therefore investigated if the changes detected in the circulating monocytes may be seen within the inflamed CNS at EAE onset. Cold exposure decreased the percentage of Ly6C^hi^ monocytes/monocyte derived cells and their MHCII expression within the CNS (Figure 5F). Furthermore, their Arg1 expression was increased, indicating a predominant anti-inflammatory M2-like phenotype (Figure 5F). Percentage and MHCII expression of microglia were similarly decreased (Figure 5G), together suggesting that they are activated to a lower extent at EAE onset under cold exposure.

Apart from monocytes and monocyte-derived cells, T cells are the main drivers of EAE pathogenicity. Cold exposure led to a decrease of pathogenic T cell cytokine signature and a reduction of GM-CSF/IL-17 and IFNg/IL-17 double producers in the CNS (Figure 5H). We detected similar changes in the dLN (Figure 5I), whereas in the mesenteric lymph nodes only GM-CSF was decreased (Figure S6F upper), and no changes were seen in the spleen (Figure S6F bottom). In the dLN, MHCII expression of Ly6C^hi^ monocytes was lower on day two (Figure 5J) but not at onset of EAE, or at steady state (data not shown). There were no consistent differences in dendritic cell populations or their activation status in the dLN at any time (Figure S6G). Collectively, these data show that cold exposure attenuates Ly6C^hi^ monocyte and T cell pathogenicity during active EAE.

### T Cell Priming by Monocytes is Critical for the Cold-Induced Attenuation of EAE

The described changes in two of the main pathogenic cell types mediating EAE provide mechanistic support for the observed amelioration of EAE by cold exposure. We therefore investigated what is the connection between monocytes and T cells during cold and EAE. Since cold exposure reduced T cell cytokine expression early during EAE and the monocyte changes were already detected under steady-state conditions, we raised the questions whether cold exposure affects the priming of T cells and what is the role of monocytes at regulating this stage. To investigate this, we used an adoptive transfer model for passive EAE induction, in which *in vitro* Th1 polarized, transgenic, MOG-specific 2D2 T cells were injected into room temperature or cold exposed mice (Figure 6A). Similar to the active EAE, cold exposure reduced MHCII expression of Ly6C^hi^ blood monocytes during the passive EAE onset after transfer of differentiated 2D2 cells, while monocyte percentages remained unchanged (Figures 6B-D and 6E). Priming T cells *in vitro* completely abolished the cold exposure-induced amelioration of EAE (Figures 6F-K upper). Similar results were obtained after adoptive transfer of Th17 differentiated cells (Figures 6F-6K bottom). These data suggest that the effects of cold exposure during the T cell priming phase are critical for the attenuation of EAE, and that cold exposure mediated MHCII downregulation may be responsible for these effects. These conclusions were supported with data from cold exposed mice that were shifted to room temperature on the day of immunization. Transferring cold exposed mice to room temperature prior to the T cell priming stage (Figure S7A) abolished the cold-induced effects on the active EAE disease (Figure S7B) and MHC expression on monocytes (Figure S7C). This is in agreement with the observation that the MHCII downregulation in monocytes under cold exposure was not sufficient to ameliorate EAE following adoptive transfer of pre-activated encephalitogenic T cells, thus making it unlikely that the observed monocyte changes are downstream mediators of effector T cells.

**Figure 6.**
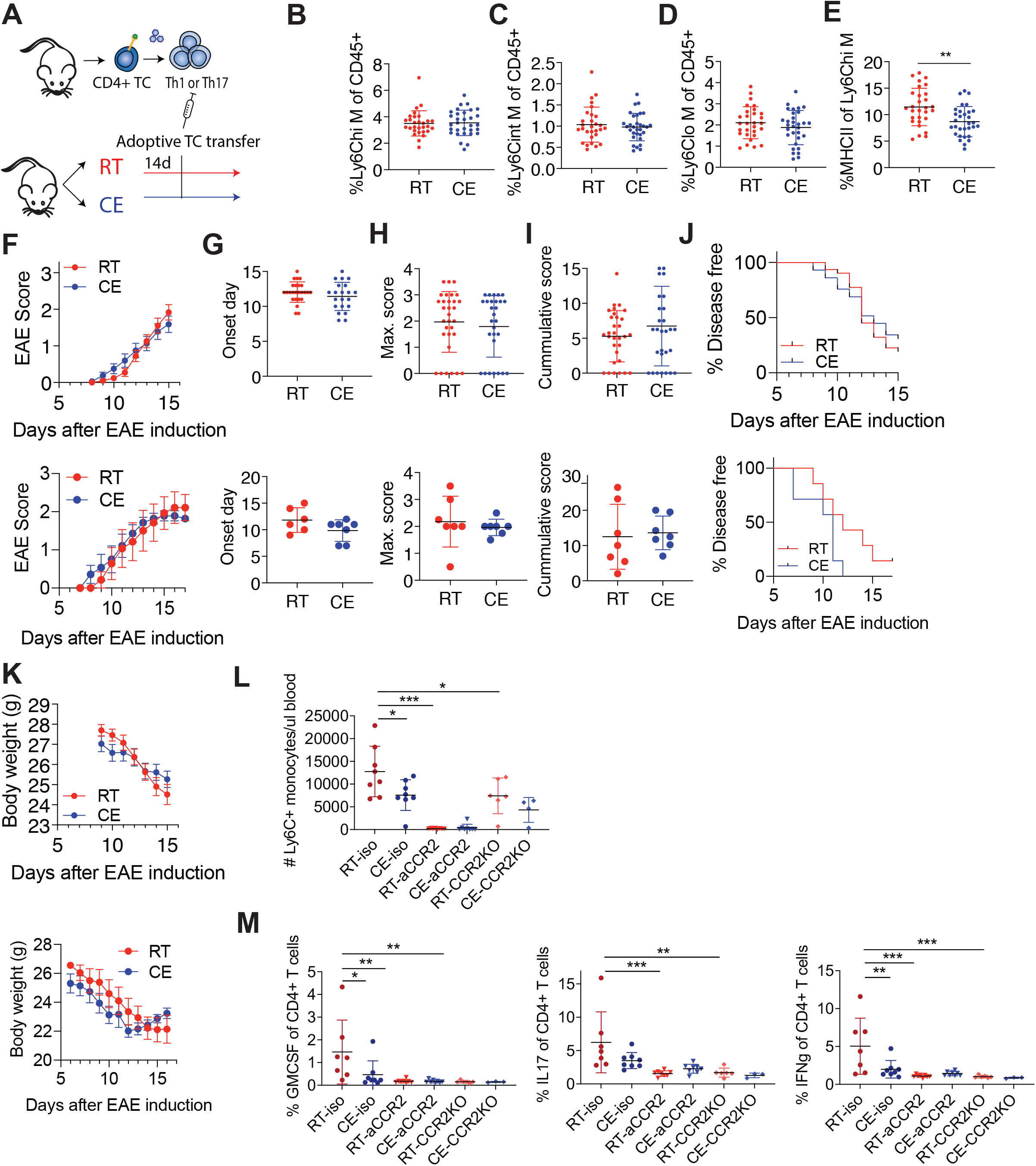
Monocyte Regulation of the T Cell Priming is Critical for Cold-Induced EAE Attenuation. (A) Scheme showing experimental setup for cold exposure and adoptive transfer EAE. T cells from transgenic 2D2 mice or actively immunized wildtype donor mice were in vitro differentiated towards Th1 or Th17 respectively and transferred into room temperature or cold exposed (2 weeks, 10°C) mice. (B-E) Flow cytometry analysis of blood cells of mice as in (A) at EAE onset. Percentage of Ly6C^hi^ (B), intermediate (C) and low monocytes of total, single, CD45^+^ cells (D). Percentage of MHCII^+^ cells of Ly6C^hi^ monocytes (E). (F-K) EAE symptoms of mice as in (A) were scored (F). Onset day of EAE disease (G), maximum score (H), cumulative score (I), percentage of disease free animals (J) and body weight (K). All panels are shown following adoptive transfer of Th1 (upper) and Th17 (lower) cells. (L-M) Isotype control antibody, CCR2 antibody injected or *Ccr2*-knockout (KO) mice were housed at room temperature or cold (10°C) for 2 weeks before and during EAE and were s.c. immunized with MOG35-55 peptide in complete Freund’s adjuvant. Two days before disease onset, monocyte depletion efficiency was analyzed in the blood (L) and at onset T cell cytokine expression in the draining lymph nodes (M) via flow cytometry. Data represents mean ± SD, significance was calculated using multiple t-test with Holm-Sidak correction, *p<0.05, **p<0.01, ***p<0.001. (B-E, F-K, L-M) Pool of 3 experiments (B-E, F-K upper panel), student’s t test with mean ± SD (B-E, G-I), two-way ANOVA with mean ± SEM (F,K) or Mantel-Cox (J). Multiple t-tests with Holm-Sidak correction with mean ±SD, ^∗^p < 0.05, ^∗∗^p < 0.01, ^∗∗∗^p < 0.001 (L-M).

To directly test whether monocytes may influence T cells during their priming phase in cold, we used two complementary approaches to deplete peripheral monocytes: anti-CCR2 (MC-21) antibody and *Ccr2*-KO mice, both of which restrict monocytes to the bone marrow (Serbina and Pamer, 2006). While anti-CCR2 completely abolished peripheral blood monocytes 2 days before EAE onset, *Ccr2*-KO mice showed significant, but incomplete loss of monocytes from the blood (Figure 6L), likely because during inflammation there could be a compensatory, CCR2-independent mechanisms for partial monocyte exit from the bone marrow. While cold exposure decreased pathogenic T cell cytokine expression in the isotype control mice, both monocyte depletion mouse models did not show any T cell cytokine expression in the dLN (Figure 6 M) irrespective of the housing temperature. Together, these data show that monocytes play a critical role in T cell priming with subsequent implications on the T cell encephalitogenic capacities.

## DISCUSSION

Our study has several fundamental implications for understanding the immune system functioning in relation to an energy-demanding metabolic stimulus, such as the change in the environmental temperature. The work provides a systematic overview of the immune cell changes in bone marrow and blood as a consequence of cold exposure during steady-state, and shows that lower environmental temperature markedly ameliorates neuroinflammation. The cold-induced gene expression alterations were accompanied by phenotypic and functional changes of monocytes. Reduced MHC class II expression and related pathways in these cells was accompanied with lowered antigen presenting capacity following various inflammatory stimuli. This cold-induced monocyte modulation resulted in reduced priming of autoreactive T cells, leading to attenuated autoimmune CNS disease. These data reveal a hitherto underestimated role of monocytes during T cell priming in autoimmunity.

Circulating Ly6C^hi^ monocytes can acquire antigen presenting functions and can exert pro- and anti-inflammatory tasks (Jakubzick et al., 2017). They therefore play an important role during infections but are also involved in autoimmune pathogenesis. The importance of monocytes and monocyte-derived cells during the effector phase of EAE is well established (Fife et al., 2000; Izikson et al., 2000; King et al., 2009; Serbina and Pamer, 2006), and in particular the relevance of migration of *Ccr2*^+^ monocytes into the CNS (Fife et al., 2000). This is in line with our observation that cold exposure reduces monocyte-derived cells during EAE in the CNS. However, our data also point out that the noted ameliorated EAE disease under cold exposure was associated with a reduced generation of encephalitogenic T cells, which was most prominent during the early phase of the disease. This suggests a cold-induced effect on the priming of naïve autoreactive CD4+ T cells, since adoptive transfer of *in vitro* pre-activated 2D2 T cells abolished the observed protection on EAE. While dendritic cells are supposed to be the primary antigen presenting cells involved in T cell priming (Ganguly et al., 2013; Steinman and Banchereau, 2007), we did not find indications that cold-exposure impacted dendritic cell function *per se*. Instead, our data rather suggest that altered monocytic function was implied in the reduced priming of autoreactive T cells under cold. Although it is known that monocytes can present antigens to T cells, their role in the context of CNS autoimmunity for T cell priming remains debated (Guilliams et al., 2018; Jakubzick et al., 2017). One previous study (Ko et al., 2014) described decreased but not abrogated IL-17 expression in *Ccr2*-DTR mice at EAE onset, and did not report differences in IFNg or GM-CSF.

However, the *Ccr2^+^* cell depletion was induced transiently and after immunization, and therefore likely did not address the initial priming phase. Another study found a minor decrease in IFNg/GM-CSF double producing CD4^+^ T cells, but no abrogation and no effect on IL-17 in *Ccr2*-KO mice before EAE onset (Ronchi et al., 2016). This mild effect on T cell pathogenicity could likely be explained by the experimental conditions, or by an incomplete monocyte depletion from the circulation in the *Ccr2*-KO mice under inflammatory conditions. Indeed, the monocyte depletion in our study was milder in the *Ccr2*-KO compared to the anti-CCR2 treated mice. We observed complete abrogation of pathogenic T cell cytokines after acute depletion of monocytes using the antibody mediated approach – effects also evident in our genetic, *Ccr2*-KO mice.

In addition to the decrease in monocyte-derived cells, cold exposure led to reduction of monocyte MHCII expression in bone marrow, blood and CNS at steady state, or during the early phase of EAE, which provides a mechanistic link to the observed reduction in T cell priming and consequently ameliorated disease course. The fact that MHCII in monocytes remained reduced even after adoptive transfer of pre-activated 2D2 cells that abrogated the protective effect of cold on EAE, strengthens the conclusions that the cold-induced monocyte changes interfere upstream of T cell effector cells. We excluded other potential mechanisms by which monocytes could affect T cell priming - neither the uptake and transport of antigen by monocytes in FITC painted mice, nor monocyte cytokine expression on transcriptional level at steady-state and EAE onset were modified under cold. Previous studies have suggested a positive feedback loop between T cells and monocytes, where T cells secrete GM-CSF to induce monocyte IL-1b expression (Croxford et al., 2015), which in return amplifies expansion of GM-CSF^+^ Th17 cells (Mufazalov et al., 2017). Our study adds an additional layer to this feedback loop, stressing that as an initial step, monocyte MHCII is critical for T cell priming.

Cold exposure induces beta adrenergic stimulation of the BAT to activate lipolysis, glucose uptake and mitochondrial biogenesis, and is extensively studied in context of obesity and metabolic disorders (Jastroch, 2017). Cold-induced BAT activation and white fat browning result in a thermogenic response that is energy demanding. As such, this metabolic response may require an energetic trade-off that competes with other energy costly programs, as for instance immune responses (Wang and Medzhitov, 2019). Several studies addressed the energy trade-off concept in the context of immune defense during infections and found that energy conserving hypometabolic states may favor the utilization of tissue tolerance as a defense against bacterial pathogens (Ganeshan et al., 2019; Weis et al., 2017). Similar to cold, caloric restriction also induces negative energy balance, reduces and modulates circulating monocytes (Jordan et al., 2019), and lowers CNS monocytes and their pathogenicity (Cignarella et al., 2018; Piccio et al., 2008). In contrast to caloric restriction that requires an intervention to reduce food intake, cold exposure ameliorates EAE despite the excessive caloric uptake. Certainly, a competition between thermogenesis and a destructive immune response would be favorable with regard to autoimmune diseases, as we see in our study. It was speculated that at low temperatures, defense of remaining thermogenesis over immunity may take place (Wang and Medzhitov, 2019). Our study is in line with this scenario and shows that cold exposure dampens autoimmunity, thus providing evidence and a conceptual advance in our understanding of the energetic trade-off at the cost of autoimmunity. In support of these conclusions, cold exposure ameliorates EAE to a higher extend in absence of UCP1 and following acclimatization and activation of the UCP1-independent thermogenic mechanisms, which function at a very high energetic cost (Ikeda et al., 2017; Reynes et al., 2019; Roesler and Kazak, 2020). The cold-induced immune changes seem to originate already in the bone marrow and can be partially provoked by beta adrenergic agonism, which may suggest contribution of the nervous system to the energy trade-off. With regard to this concept, it is important to stress that while evidently protective against autoimmunity, cold exposure increases susceptibility to certain viral infections (Foxman et al., 2015; Jaakkola et al., 2014). Thus, our work could be relevant not only for CNS autoimmunity, but also other immune-mediated or infectious diseases, which warrants further investigations. Our study reveals that cold exposure orchestrates immunologic reprogramming that protects from CNS autoimmunity, showing that monocytes and the monocyte MHCII expression are crucially implicated during T cell priming in the course of an autoimmune response. These findings provide critical insights into the pathogenic mechanisms of neuroinflammation, and could lead to future preventive and therapeutic approaches for MS and other autoimmune diseases.

## ACKNOWLEDGEMENTS

We thank the iGE3 Genomics Platform at the University of Geneva for mRNA sequencing and mapping reads, the Flow Cytometry core facility at the CMU of the University of Geneva for help with flow cytometry, Christelle Veyrat-Duberex, Florian Visentin, Melis Colakoglu, Giovanni di Liberto, Salvatore Fabbiano and Karim Hammad for experimental assistance, Sergei Startchik for the help on automatized lipid droplet quantification, and Stephanie Hughes and Pierre Guermonprez for organizing and providing B16-GMCSF cells, respectively, Thomas Korn for the Th17 EAE protocol and Claes Wollheim for discussions. This work was supported by the CONFIRM grant of the Hôpitaux Universitaires de Genève (HUG) Foundation (no. RC2-09) and the Swiss Multiple Sclerosis Society Grant to DM and MT; the European Research Council (ERC) under the European Union’s Horizon 2020 research and innovation programme (ERC Consolidator Grant agreement No. 815962) to MT; Swiss National Science Foundation (SNSF) grant to DM (310030_173010); and SNSF Professorship grants (PP00P3_144886 and PP00P3_172906) to MT.

## AUTHOR CONTRIBUTIONS

M.S. set up and performed experiments, analyzed and interpreted data, and drafted figures and manuscript. K.S. set up and performed experiments, and analyzed and interpreted data. N.H. and M.S. performed RNA-seq analysis. D.R., N.S.Z., M.A.R., C.C., I.V., N.P., B.K. and O.S. helped with experiments. I.W. assisted in histological processing. M.G. assisted in setup of and D.C. performed and analyzed FACSymphony. M.K. developed and applied computer-assisted image analysis algorithms. MM provided the MC-21 antibody. D.M. and M.T. conceptualized, designed and supervised the project, interpreted data, and wrote the manuscript.

## Supplemental Figure Legends

**Figure S1. (Related to Figure 1).**
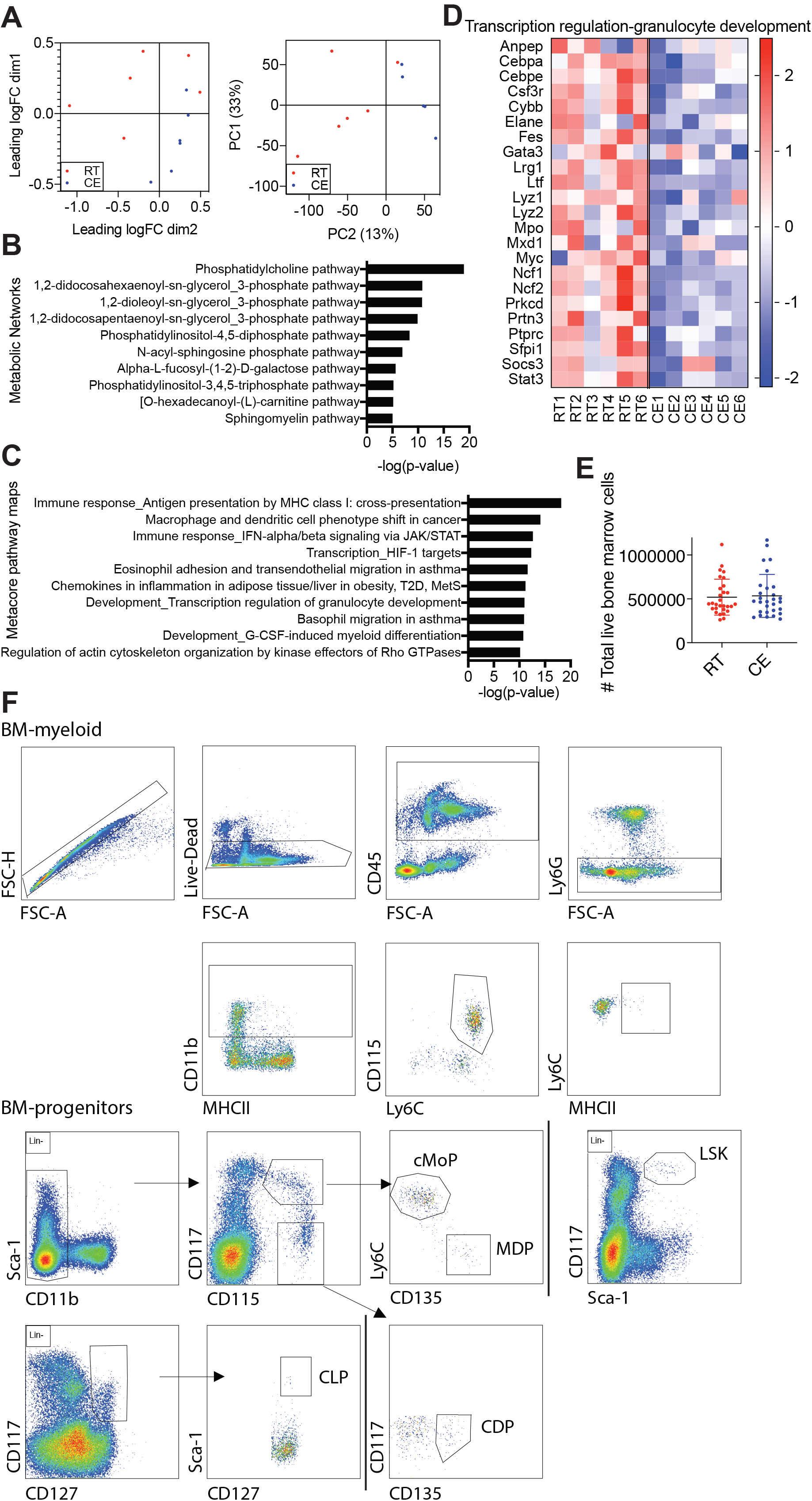
(A) Multidimensional scaling and principal component analysis of the bone marrow from mice after 2 weeks of cold exposure at 10°C compared to room temperature mice. (B) 10 most enriched Metacore Metabolic Networks of mice as in (A). (C) Metacore Pathway Maps analysis identifying the 10 most differentially regulated pathways of mice as in (A). (D) 1 of 4 heatmaps that were consistently changed (> 80% of genes in the same direction) from the top 10 deregulated Metacore Pathway Maps of mice as in (A). (E) Absolute lymphocyte numbers within the bone marrow of mice as in (A) as determined by flow cytometry. Shown is a pool of 5 experiments, mean ±SD. (F) Bone marrow monocytes were gated as live, single, CD45^+^, Ly6G^-^, CD11b^+^, CD115^+^, Ly6C^+^ cells and MHCII^+^ population determined. Bone marrow progenitors were all gated as single, live, CD45+, Lin-(NK1.1, CD3, CD19, Ter119, Ly6G) and CD11b^-^, Sca1^+^, CD117^hi^, CD115^+^, Ly6C^hi^ (cMoP), CD11b^-^, Sca1^+^, CD117^hi^, CD115^+^, Ly6C^-^ (MDP), CD11b^-^, Sca1^+^, CD117^-^, CD115^+^, CD135^+^ (MDP), CD117^+^, CD127+, Sca-1^+^ (CLP), Sca-1^+^, CD117^+^ (LSK). (C-D) The cutoff on differentially regulated genes considered for the pathway analysis is p<0.05 and pathways are considered deregulated with p<0.05. Shown are –log(p-value).

**Figure S2. (Related to Figure 2).**
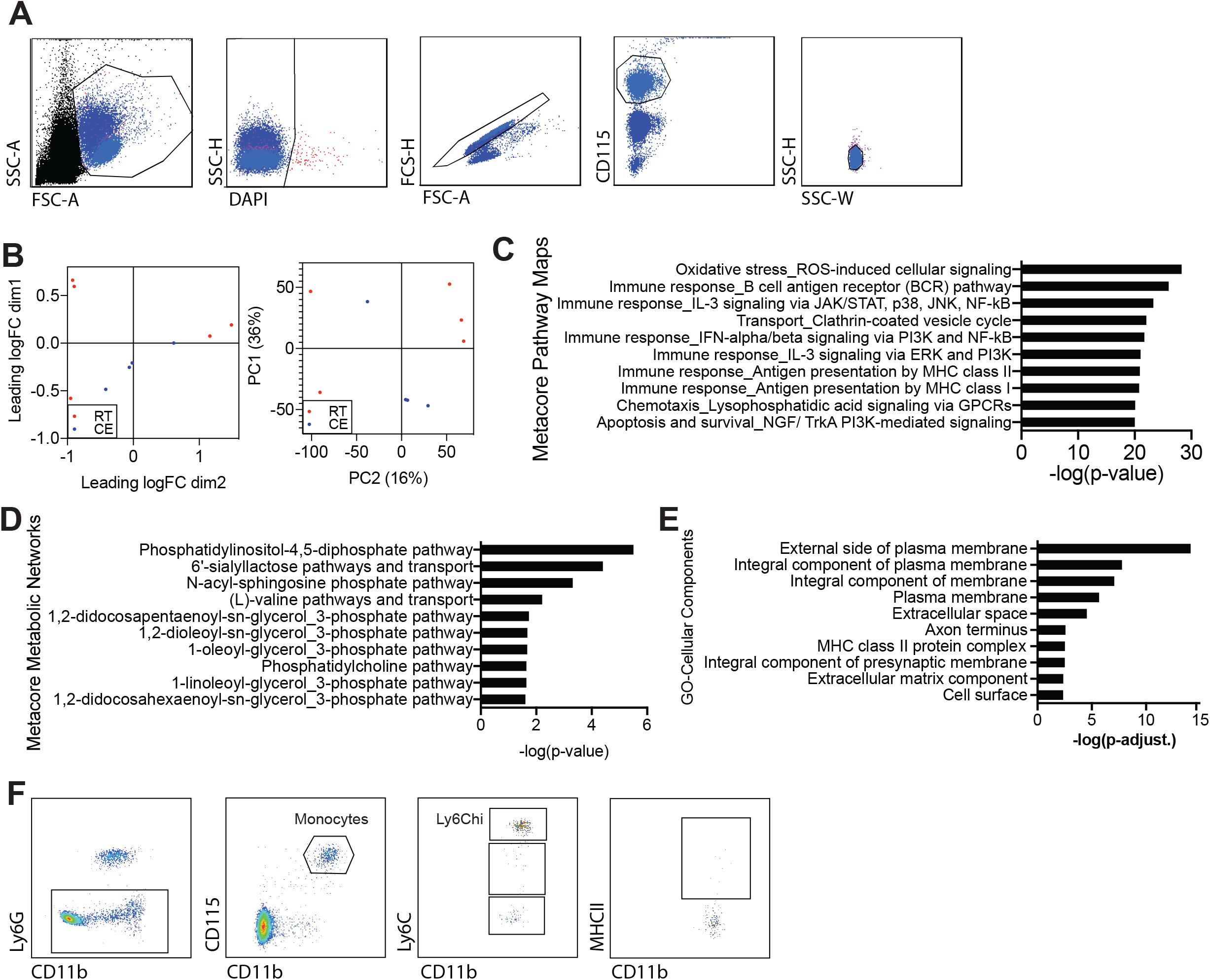
(A) Gating strategy for the FACS (CD115-PE+) sort after MACS (anti-PE) sort before RNA sequencing from blood of 2 weeks cold exposed (10°C) compared to room temperature mice. Sorted were DAPI-, single, CD115+ cells. (B) Multidimensional scaling (left) and principal component analysis (right) after RNA sequencing of monocytes as in (A). (C-E) 10 most differentially regulated Metacore Pathway Maps without p-value cutoff on considered genes (C), Metacore Metabolic Networks (D) and Gene Ontology Cellular Components (E). Genes were considered as differentially regulated genes with p<0.05 and pathways are considered deregulated or enriched with p<0.05. Shown are –log(p-value). (F) For flow cytometry analysis, blood monocytes were gated as Ly6G^-^, CD11b^+^, CD115^+^, Ly6C^hi^/^int^/^lo^ and MHCII^+^ positive population was gated.

**Figure S3. (Related to Figure 3).**
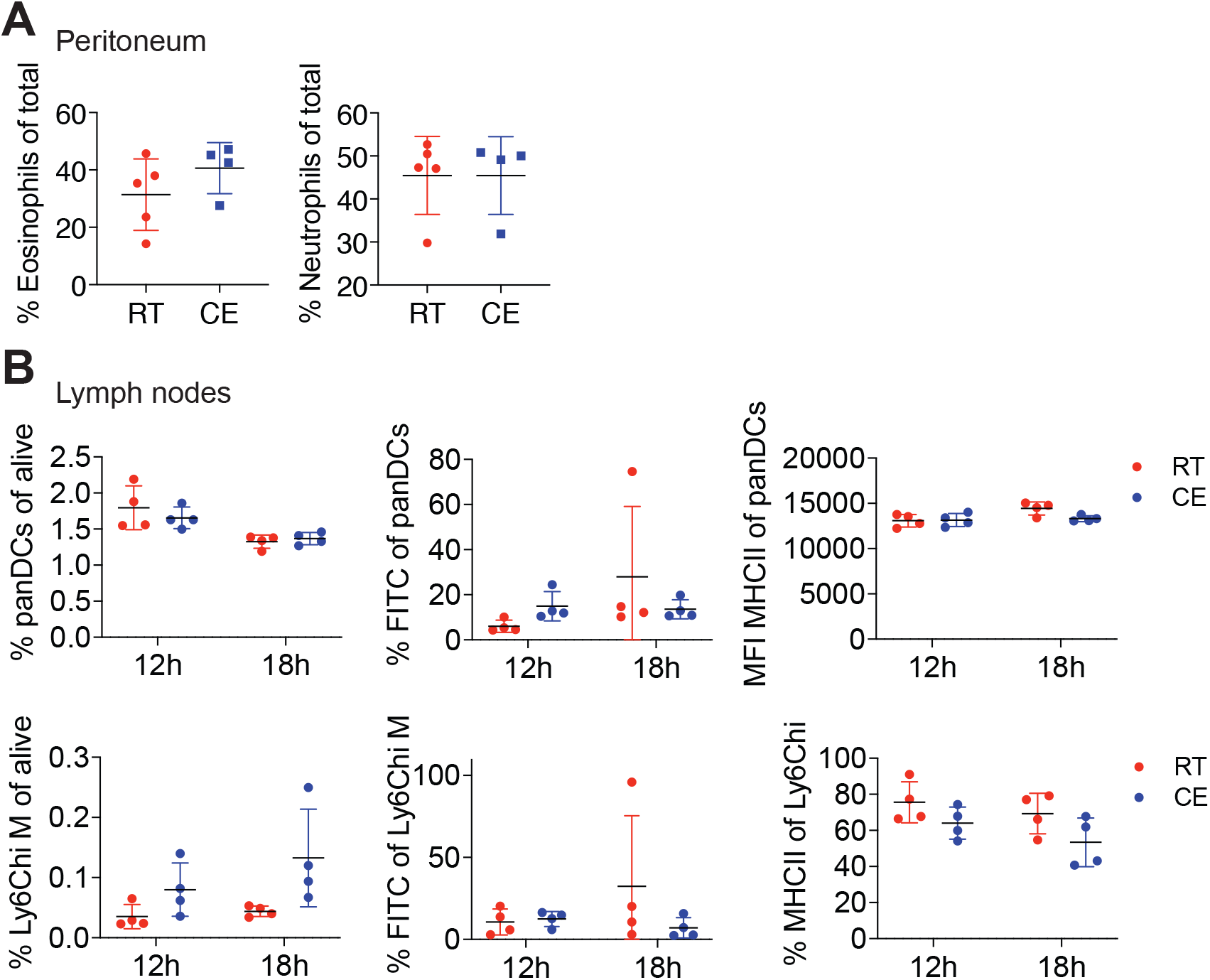
(A) Peritoneal fluid cells were analyzed by flow cytometry 24h after i.p. injection with thioglycollate into two weeks cold exposed (10°C) or room temperature mice. Shown is mean ±SD. (B) Flow cytometry analysis of lymph node draining cells 12 and 18 hours after FITC skin painting on one flank of 2 weeks cold exposed (10°C) or room temperature mice. Percentage of panDCs (upper) and Ly6C^hi^ monocytes (bottom), their FITC uptake and MHCII expression. Shown is mean ±SD, two-way ANOVA. Every dot represents one animal and the experiment is representative of two.

**Figure S4. (Related to Figure 4).**
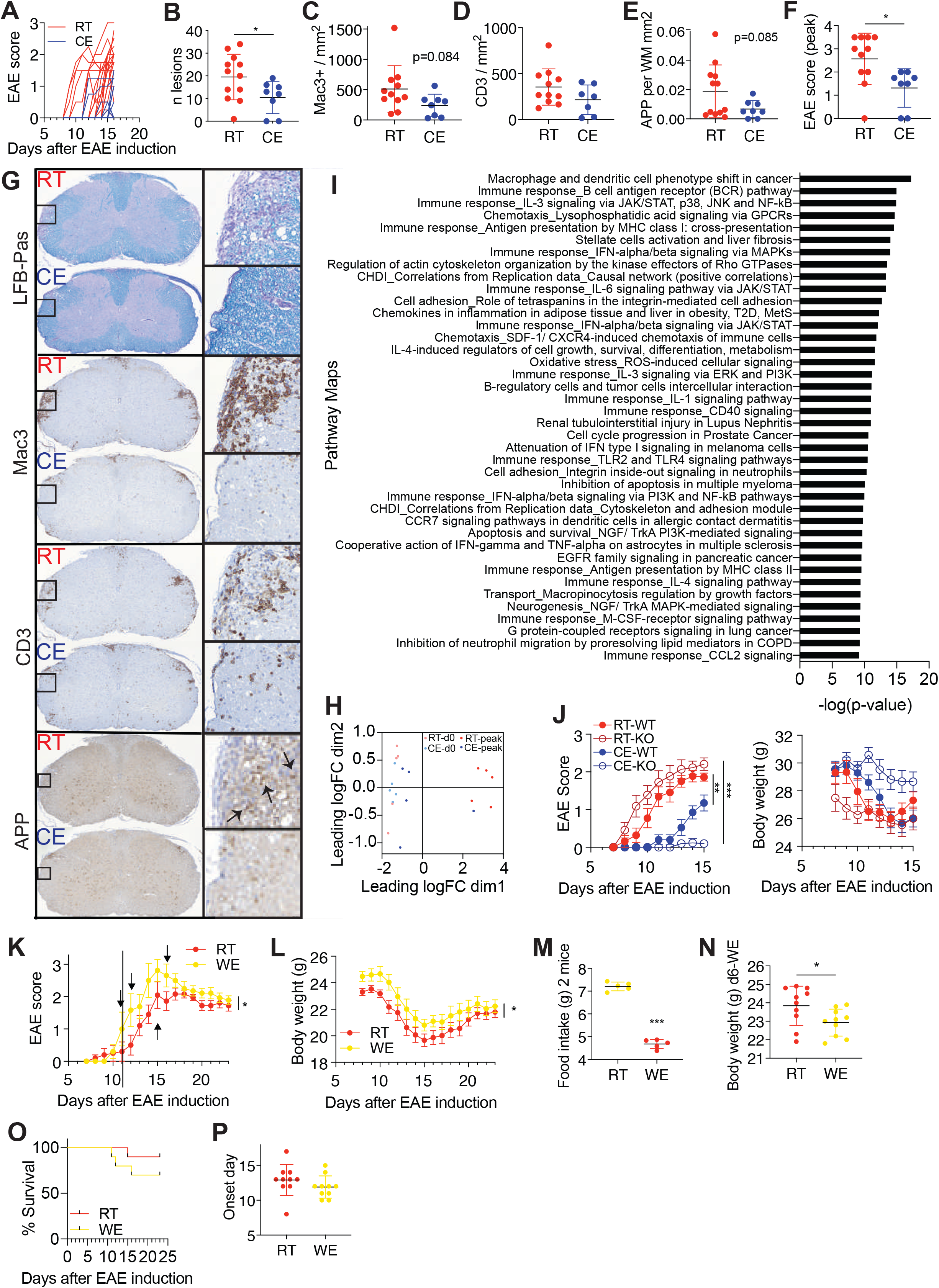
(A) Individual EAE (as in Figure 4A) curve of each room temperature and cold exposed mouse. (B-G) Histology on sagittal spinal cord sections of cold exposed and room temperature peak EAE mice. Number of lesions that were found in both LFB-PAS and APP staining were quantified in APP staining (B). Immunohistochemistry using anti-mac3 (C) and anti-CD3 antibody (D). APP staining was quantified within areas identified as demyelinated in LFB-Pas staining and presented as percentage per white matter area (E). EAE disease score at sacrifice of mice in which histology was performed (F). LFB-Pas stain shows demyelinated areas within the white matter in purple (first panel), anti-mac3 stains mac3+ macrophages (second panel), anti-CD3 stains T cells in brown (third panel) and amyloid precursor protein (APP) accumulates at axons with impaired axonal transport, a surrogate for acute axonal damage (fourth panel). Arrows identify punctuated APP-positive axonal spheroids. (G). Anti-mac3, anti-CD3 and anti-APP stainings were developed with DAB and nuclei counterstained with hemalum. (H, I) Multidimensional scaling analysis of spinal cord from mice as in Figure 4J (H). Top 35 Metacore Metabolic Network pathways of mice with EAE, genes with p<0.05 were included for analysis (I). (J-O) EAE disease curve of warm exposed (34°C, started one week before immunization) or room temperature mice. Arrows indicate individual mice that died or had to be sacrificed, which received a score of five on the day of death (J). Body weight during the course of EAE (K). Food intake of two mice per cage within 24 h on day 6 of warm exposure (L) and body weight on day six of warm exposure (M), before immunization. Survival of mice during EAE (N) and day of EAE onset (O). (B-F, J-O) Shown is mean ±SEM, two-way ANOVA (J, K); Mantel-Cox (N), student’s t test (B-F, L-M, O) with mean ±SD, ^∗^p < 0.05, ^∗∗^p < 0.01, ^∗∗∗^p < 0.001.

**Figure S5. (Related to Figure 5).**
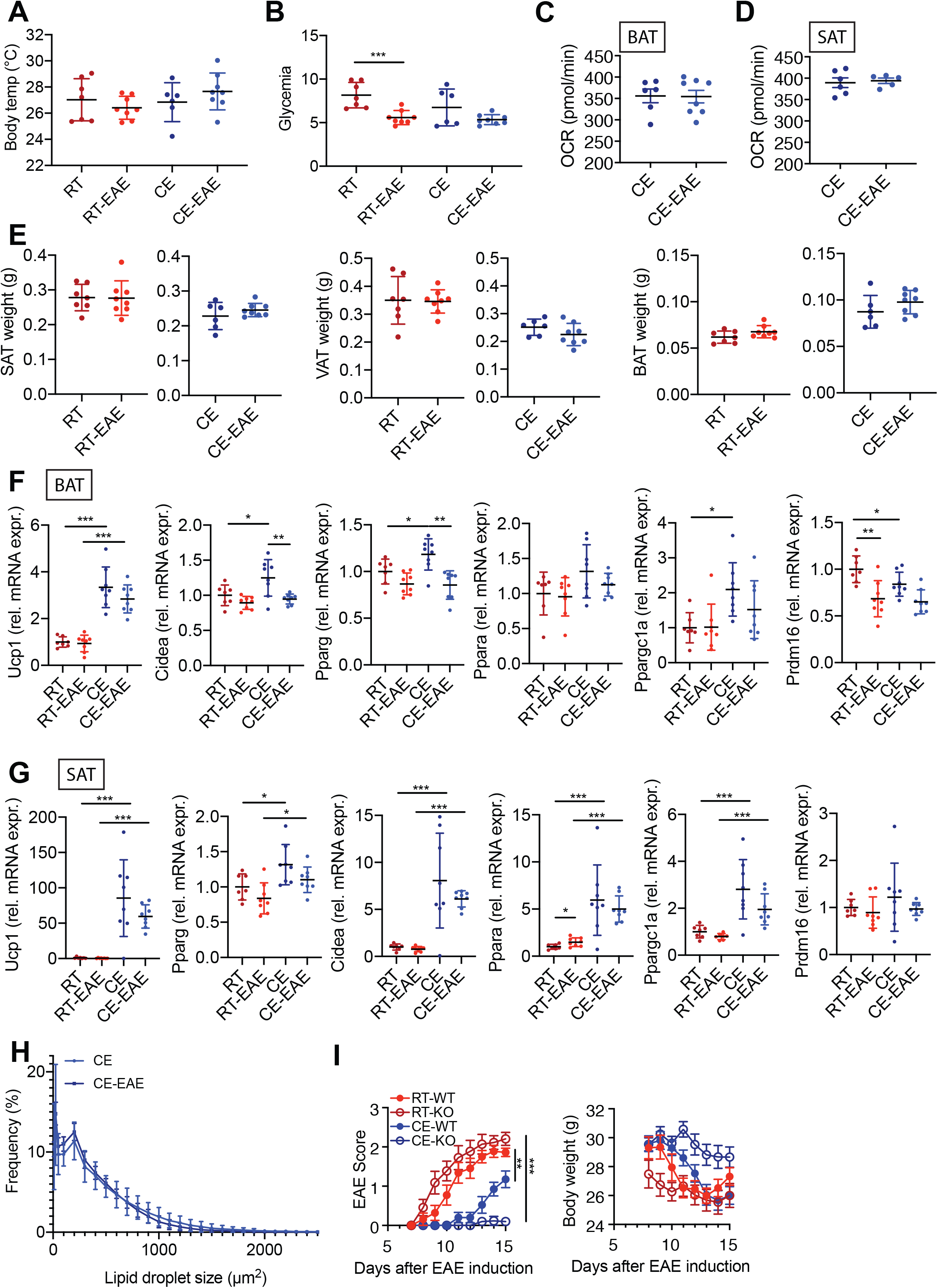
(A) Body temperature shown is the mean temperature of the two eyes as determined via infrared images of mice on day 7 of EAE and steady state. Mice were cold exposed for 2 weeks before and during EAE or steady state. Control mice were at room temperature for the same duration. (B) Glycemia of mice as in (A) on day 8 of EAE after 3 hours of fasting. (C-D) Oxygen consumption rate of interscapular brown (BAT, C) and inguinal subcutaneous adipose tissue (SAT, D) on day 8 of EAE or in steady state after 3 weeks of cold exposure (CE) and 3 hours of fasting. Technical replicates were used for BAT. (E) Adipose tissue weights of interscapular brown (BAT, left), inguinal subcutaneous adipose tissue (SAT, middle) and perigonadal visceral adipose tissue (VAT, right) of mice as in (C). (F-G) mRNA expression of browning markers relative to *B2m* and *36b4* in interscapular brown (BAT, F) and inguinal subcutaneous adipose tissue (SAT, G) of mice as in (B) as determined by qPCR. (H) Quantification of lipid droplet size distribution in ingSAT. Slides were analyzed in technical duplicates from different layers of the tissue and averaged. (I) Symptoms of EAE (left) and body weight (right) in *Ucp1*-knockout (KO) and their respective wildtype (WT) littermate controls housed at room temperature or cold for 1 week before and during EAE. Shown is 1 representative (n=10-15 per group) of 2 experiments. (A-G, I) Multiple t-test with Holm-Sidak correction (A-B, F-G), student t-test (C, D), mean ±SD, two-way ANOVA with mean ±SEM (I), ∗p < 0.05, ∗∗p < 0.01, ∗∗∗p < 0.001.

**Figure S6. (Related to Figure 6).**
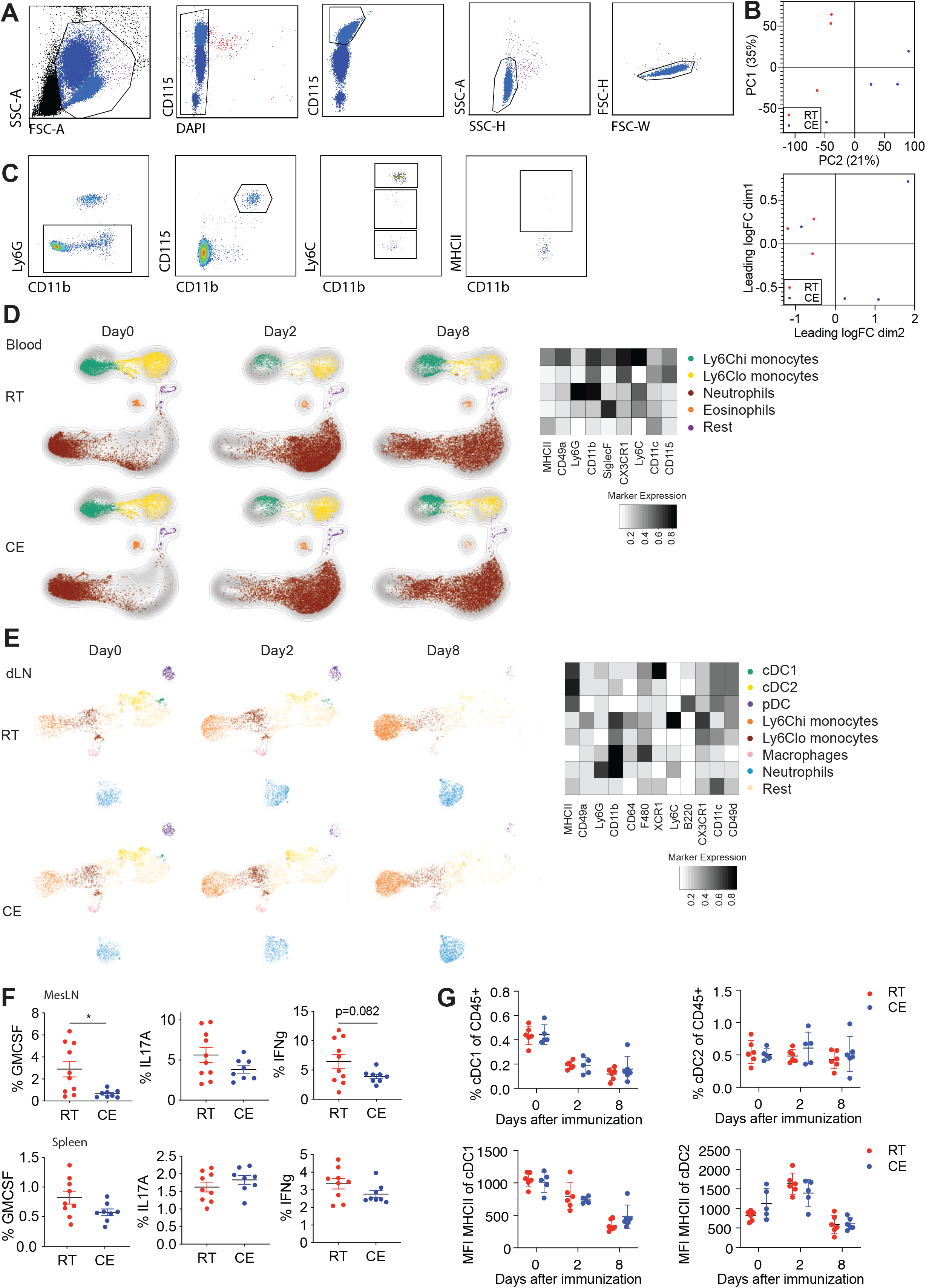
(A) Gating strategy for the FACS sort after MACS sorting before RNA sequencing (as in Figure 6A). Sorted were DAPI-, CD115^+^, single cells. (B) Multidimensional scaling and principal component analysis of FACS sorted blood monocytes at EAE onset as in Figure 6A. (C) Gating strategy for MHCII expression on Ly6C^hi^ blood monocytes of mice as in Figure 6A. Cells were gated as single, Ly6G^-^, CD115^+^CD11b^+^, Ly6C^hi^, MHCII^+^. (D-E) Blood (D) and lymph node (E) immune cells of mice as in Figure 6A on different days after immunization were visualized using UMAP and clustered using the FlowSOM algorithm in R. Heatmap shows median relative expression of all panel markers. (F) Flow cytometry analysis of mesenteric LN (upper panel) or spleen cells (lower panel) of mice as in Figure 6A at EAE onset. Percentage of cytokine expression in CD4^+^ T cells. Shown is mean ±SD, significance was calculated using student’s t test, * p<0.05. (G) Flow cytometry analysis of lymph node dendritic cells and their MFI of MHCII on different days after immunization.

**Figure S7. (Related to Figure 6).**
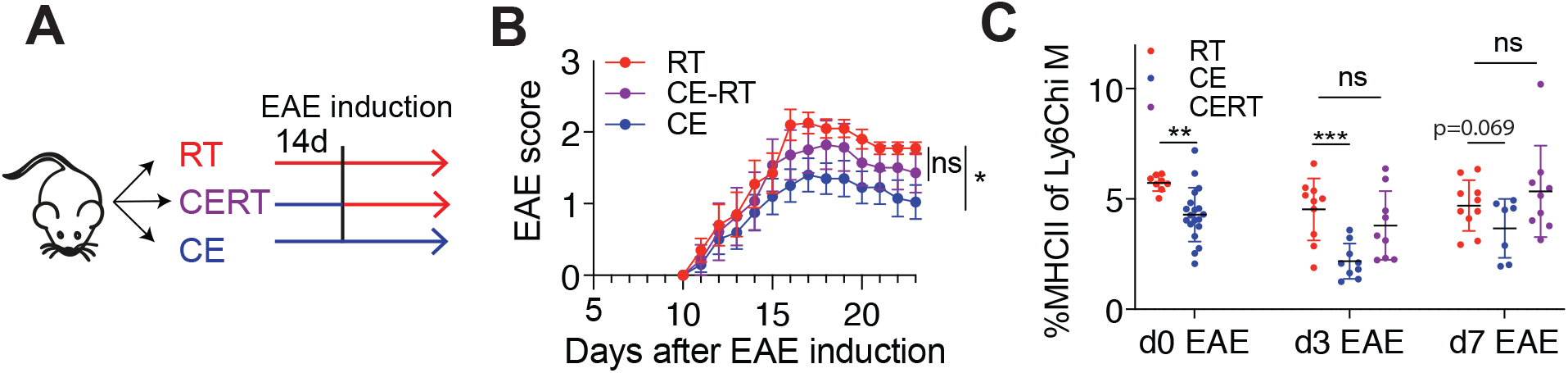
(A-B) Scheme showing experimental setup for cold exposure and EAE (A). Cold exposure either started two weeks before and continued during EAE (blue, CE) or switched to room temperature from the day of immunization (violet, CE-RT) and compared to room temperature mice (red, B). EAE disease curve shown as mean ±SEM, two-way ANOVA, ^∗^p < 0.05. (C) Percentage of MHCII expression of Ly6C^hi^ blood monocytes via flow cytometry on day zero, day three and day seven after EAE induction. Multiple t-test with Holm-Sidak correction with mean ±SD, ^∗^p < 0.05, ^∗∗^p < 0.01, ^∗∗∗^p < 0.001.

## METHODS

### RESOURCES TABLE

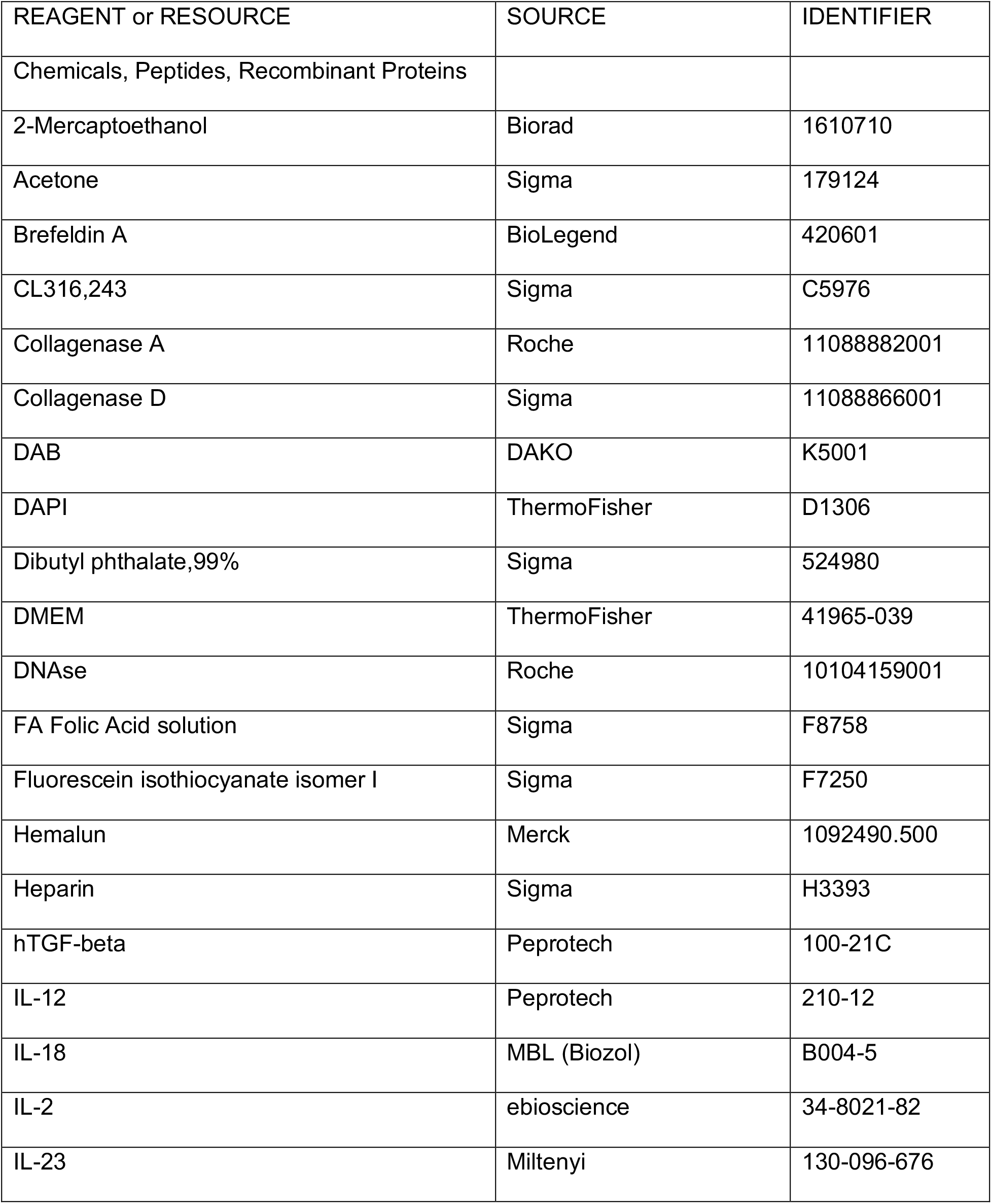
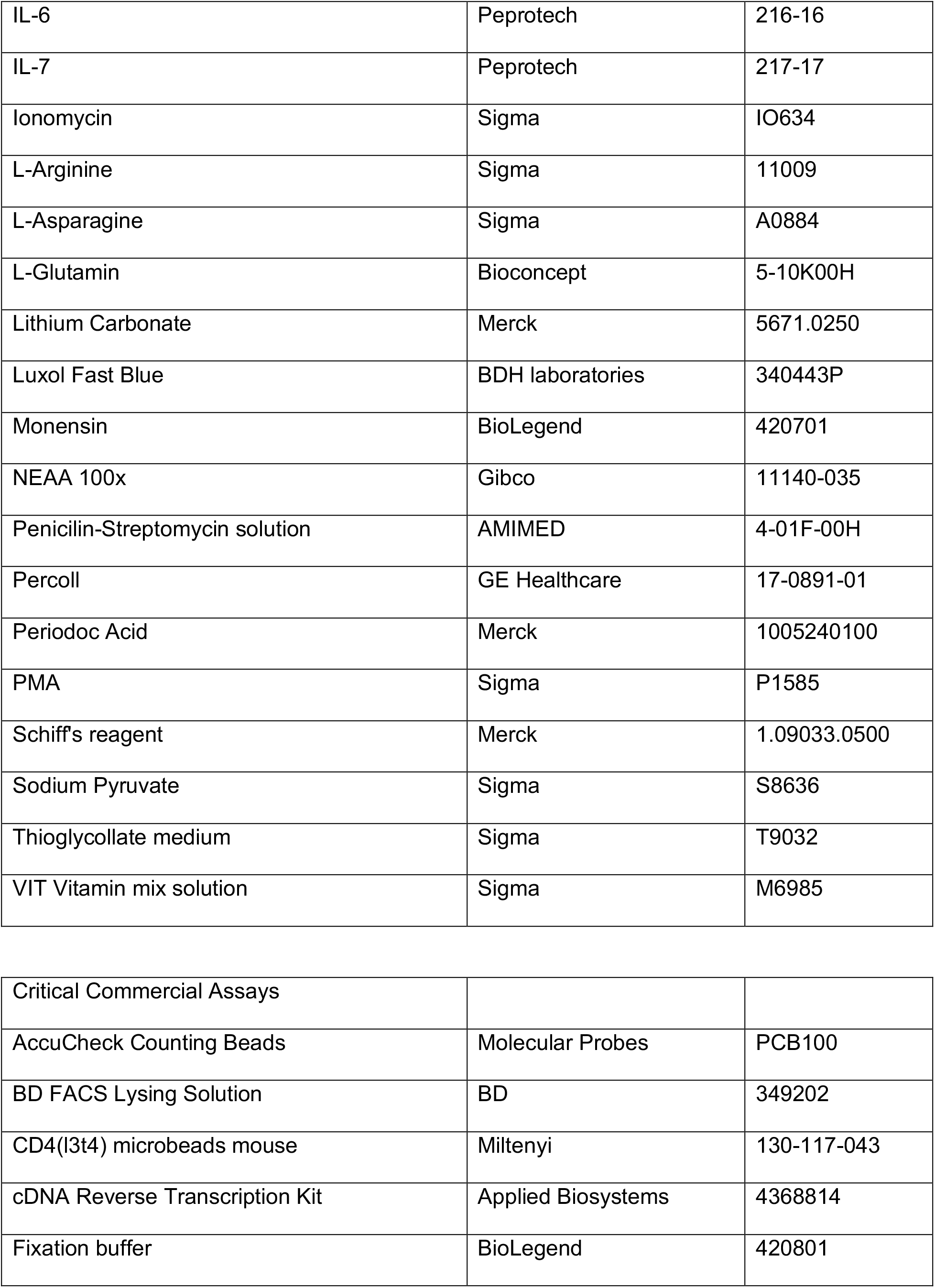
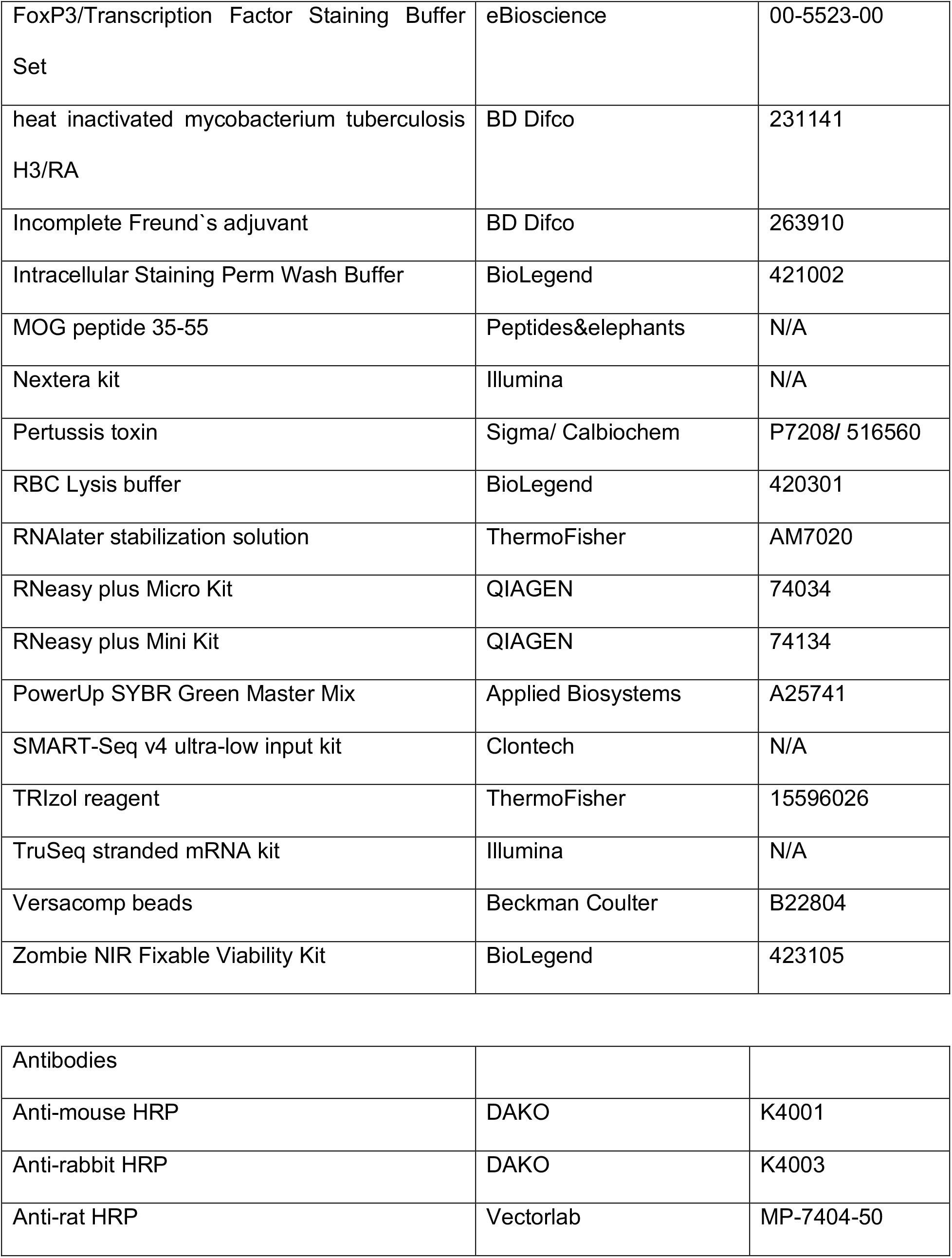
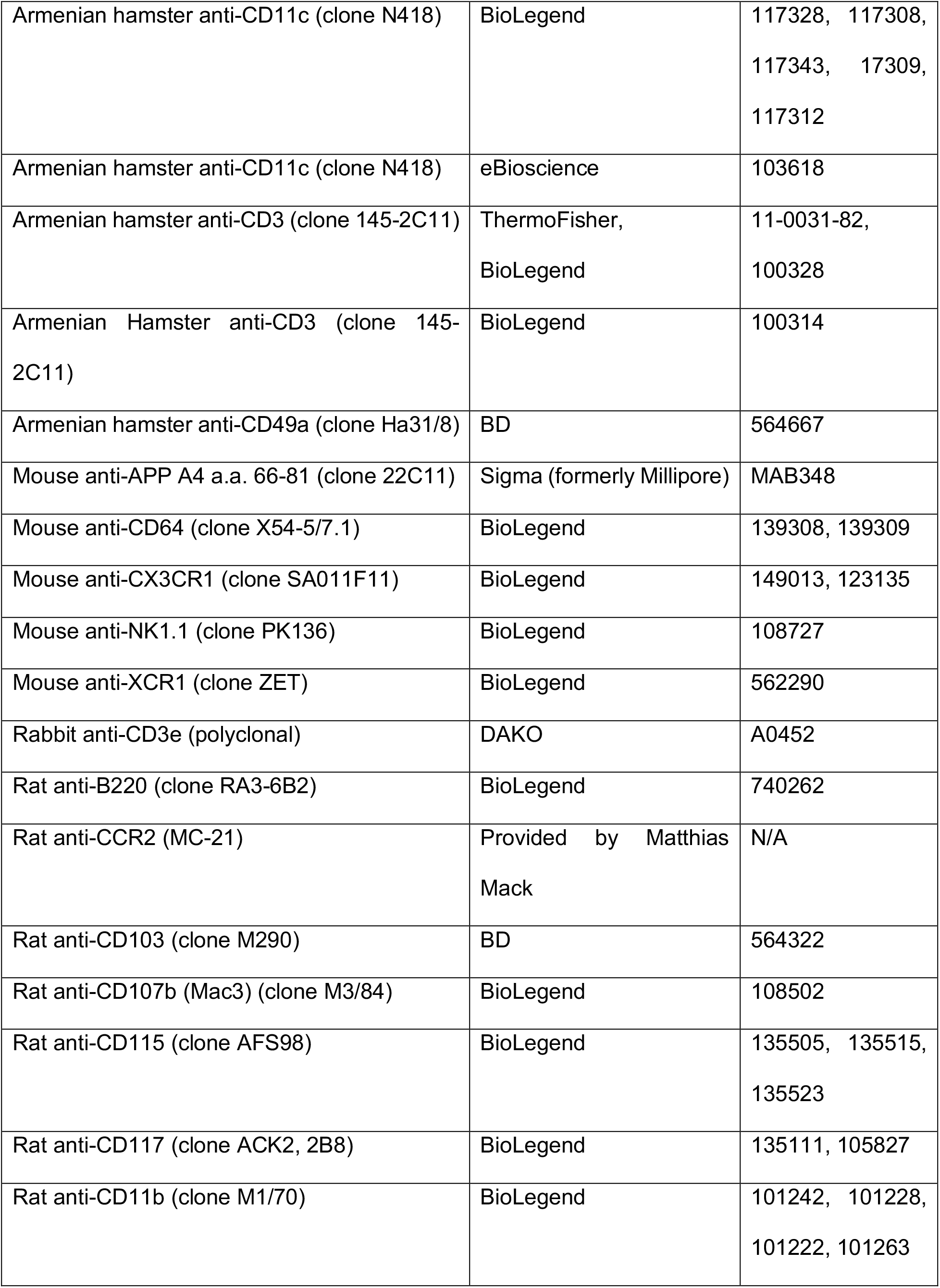
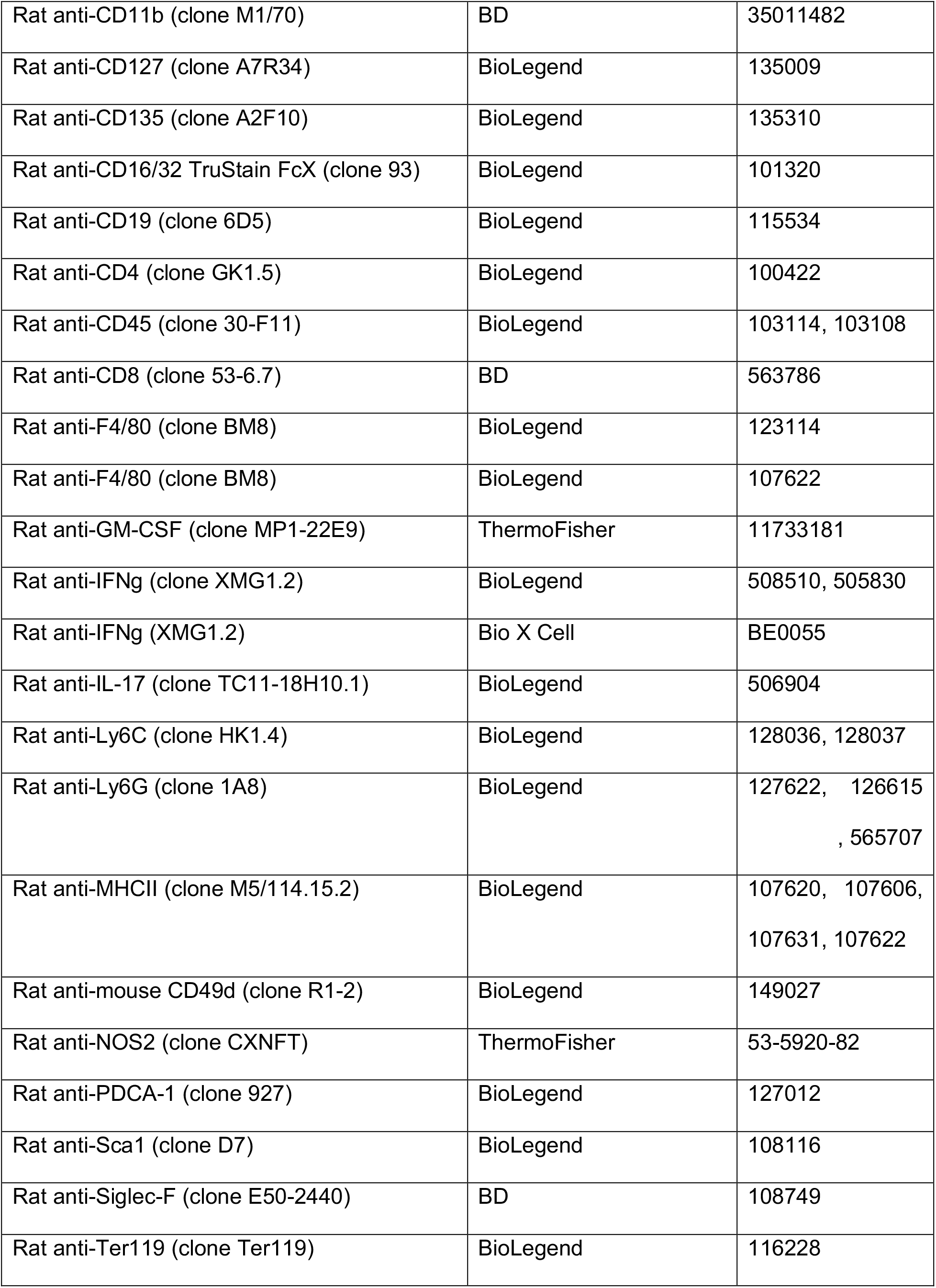
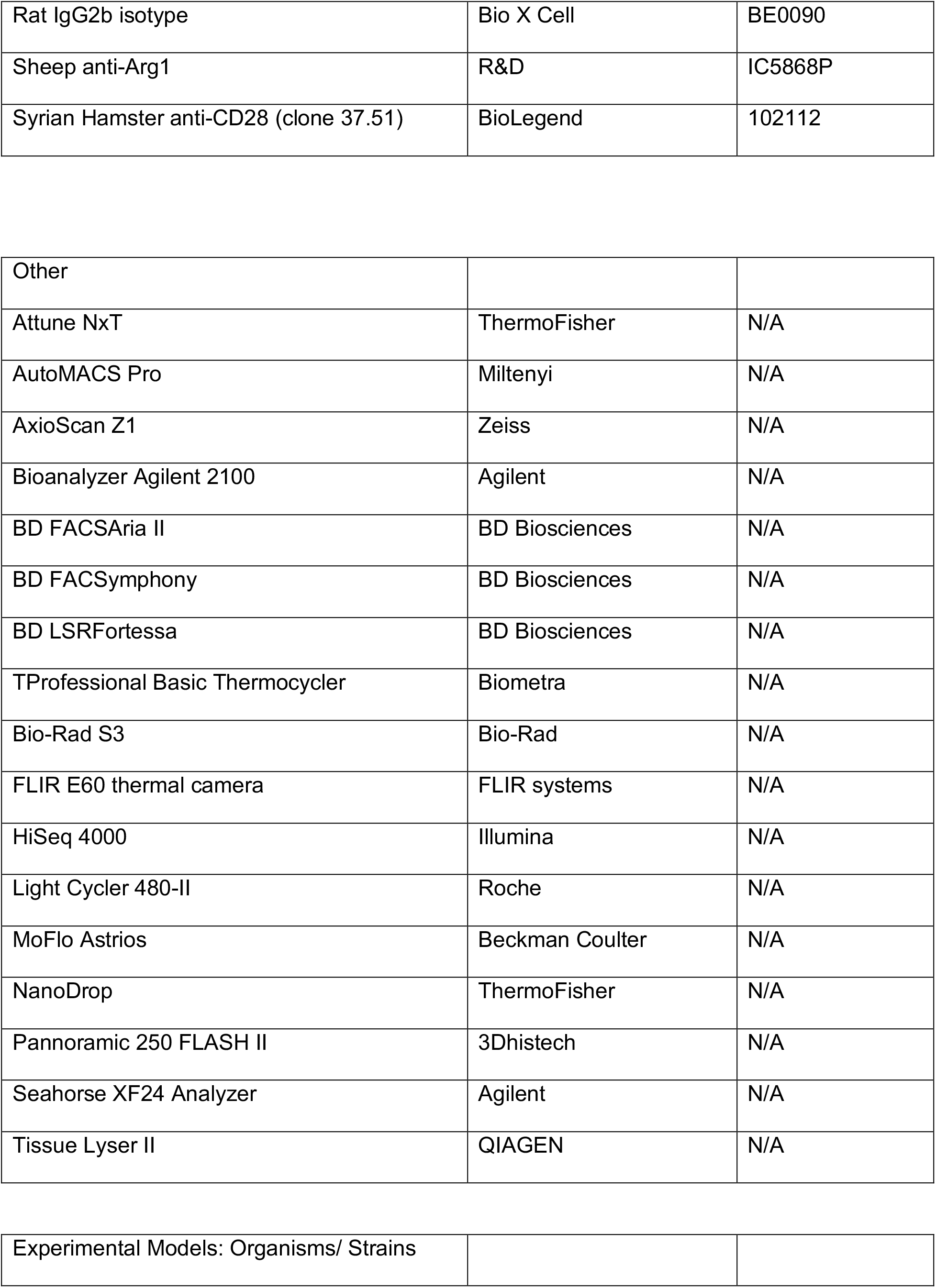
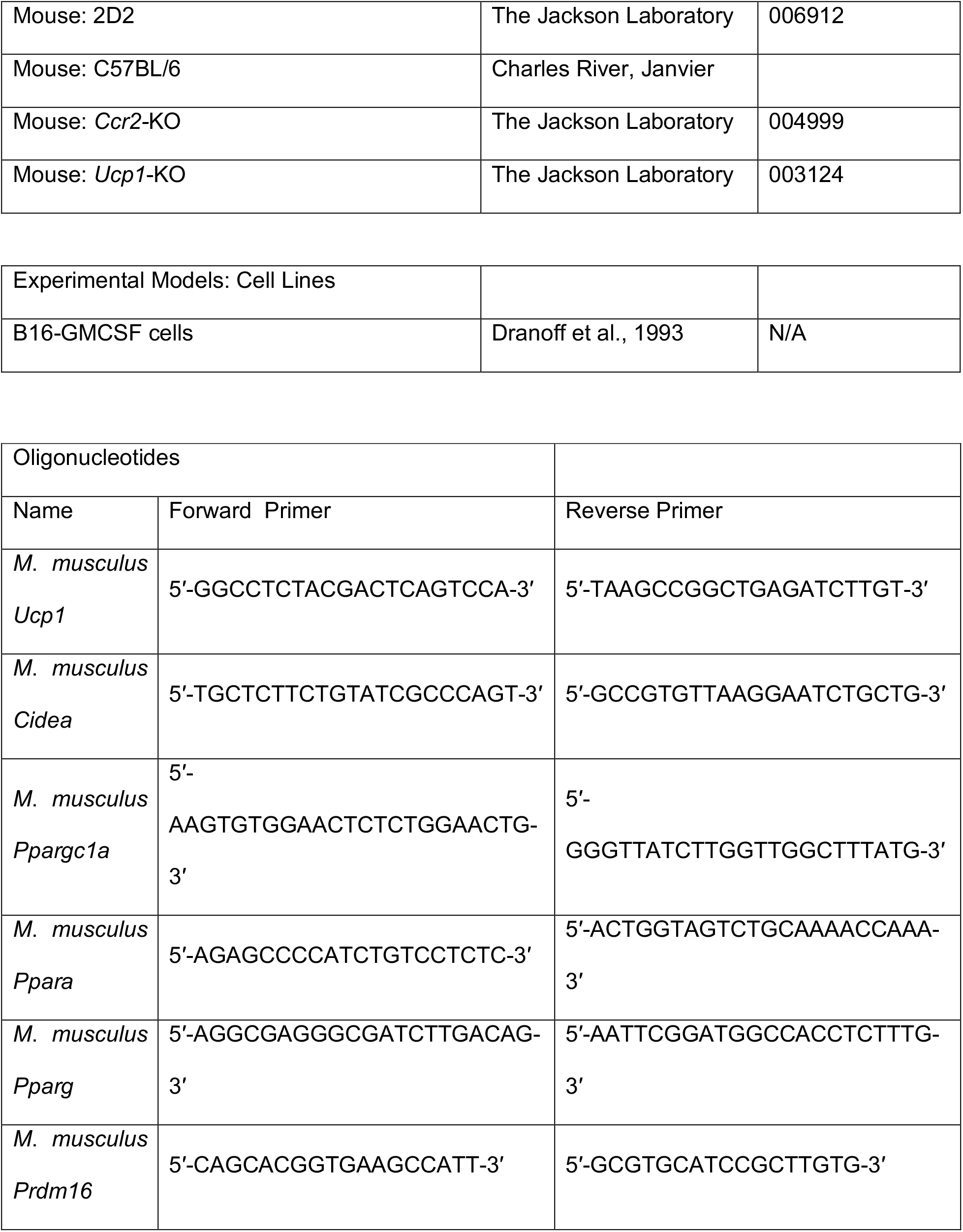
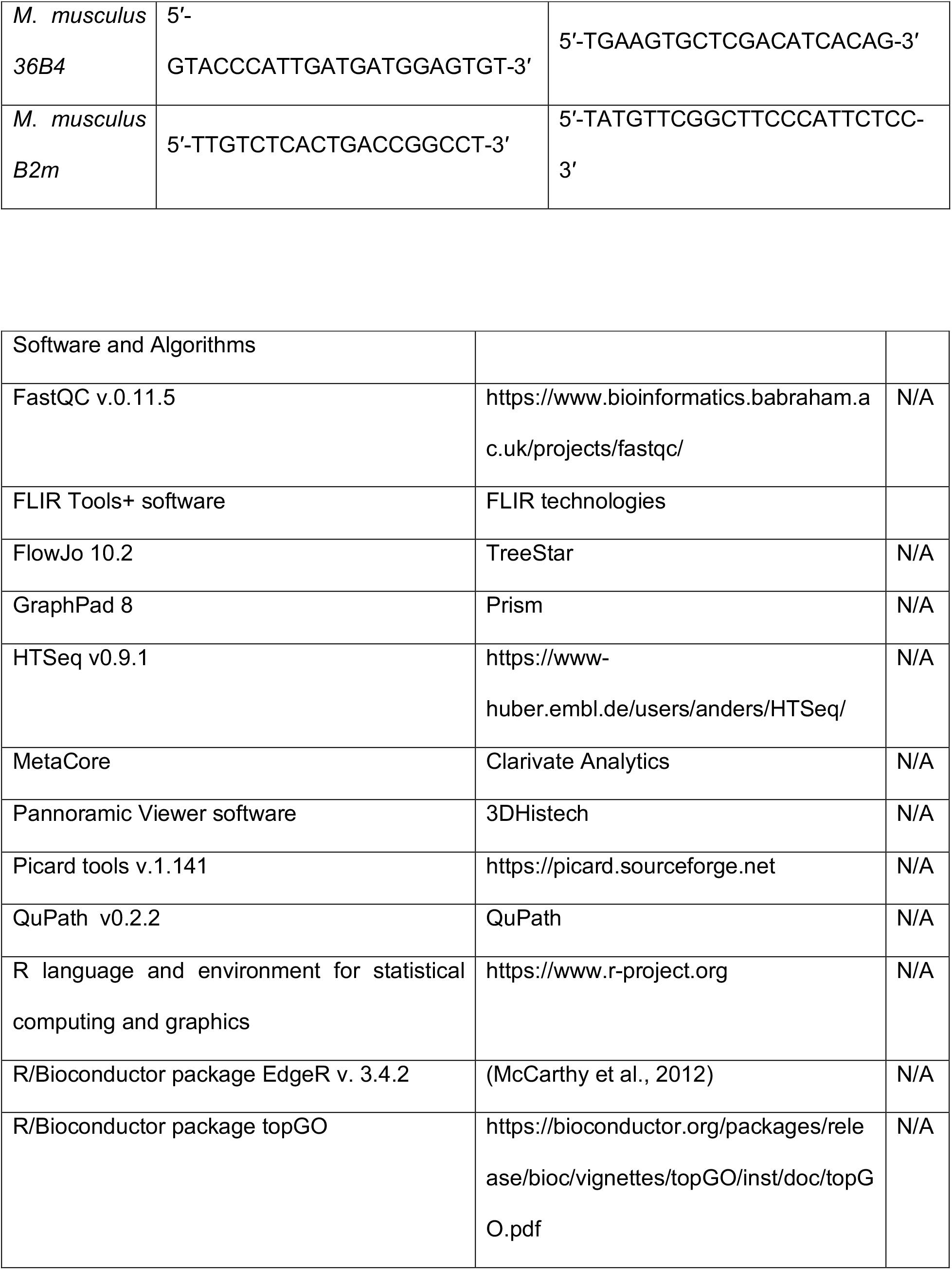
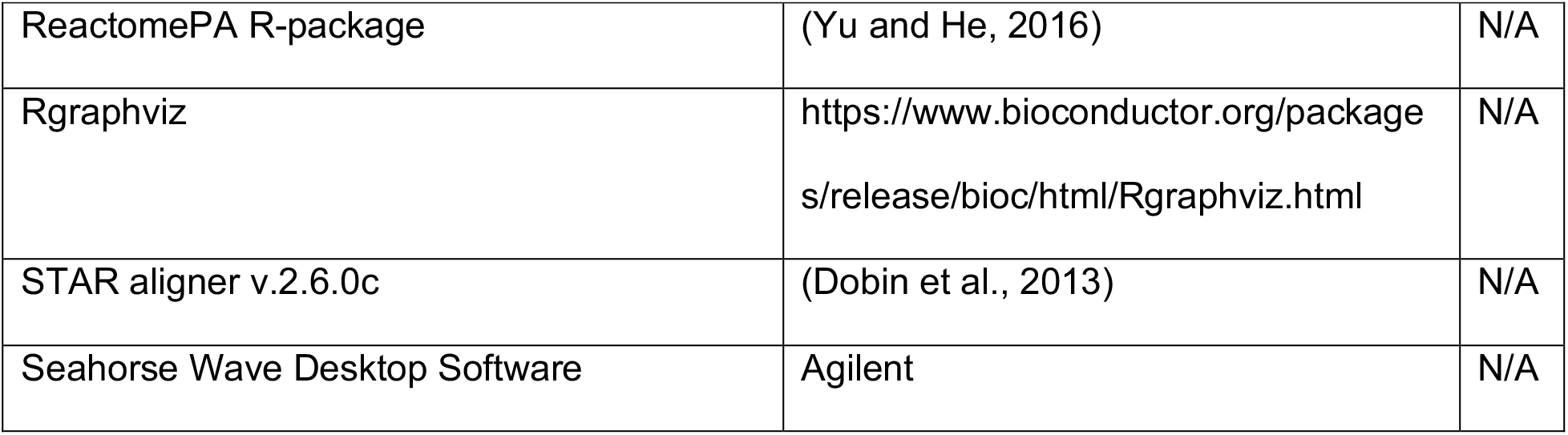

## EXPERIMENTAL MODEL AND SUBJECT DETAILS

### Mice

Male 8 weeks old C57BL/6J mice were obtained from Charles River, or Janvier (France). *Ucp1*-KO (The Jackson Laboratory), their littermate controls, *Ccr2*-KO (The Jackson Laboratory) and 2D2 (The Jackson Laboratory) mice were bred in house. Mice were housed in a specific pathogen free (SPF) facility in 12 h day/night cycles, fed irradiated standard chow diet and given water from autoclaved bottles. Cold exposure was applied in a light- and humidity-controlled climatic chamber (TSE, Germany) under SPF conditions, at 10 °C for 2 weeks before immunization with an additional initial acclimatization period of 5 days at 18°C and 5 days at 14°C, as well as during EAE. Warm exposure of 34°C was applied for 7 days before immunization, and during EAE. All climate chamber experiments were performed with 2 mice per cage without bedding and nesting material for enrichment to ensure controlled temperature conditions. CL316,243 (Sigma) was i.p. injected daily 7 days before immunization and until sacrifice. Body temperature was determined with infrared camera FLIR E60 (FLIR, UK) taking the mean temperature of both eyes using the FLIR Tools+ software. All animal experiments were performed at the University of Geneva with authorization by the responsible Cantonal authority and in accordance with the Swiss law for animal protection.

## METHODS DETAILS

### EAE induction and scoring

For active EAE immunization, complete Freund’s adjuvant was prepared by mixing 10mg/ml heat inactivated mycobacterium tuberculosis H3/RA (Difco) with incomplete Freund’s adjuvant (Difco). To create an emulsion, this was mixed 1:1 with MOG peptide 35-55 (4 mg/ml, peptides & elephants). Mice were anesthetized and 50 ul of the emulsion was injected into the left and right flank each (200 ug peptide/mouse), followed by i.v. injection of 300 or 400 ng pertussis toxin (Sigma or Calbiochem). After two days, mice received additional 300 or 400 ng pertussis toxin i.p.

For Th1 adoptive transfer EAE, splenocytes of 2D2 T cell receptor transgenic mice were cultured for 2 days with MOG35-55 peptide (20 µg/ml), IL-2 (5 ng/ml, ebioscience) and IL-7 (5 ng/ml, Peprotech), 4 days with IL-2 (5 ng/ml) and IL-7 (5 ng/ml) and 1 day with anti-CD3/28 (0.5 ng/ul, Biolegend), IL-12 (20 ng/ml, Peprotech) and IL-18 (25 ng/ml, MBL (Biozol)) at 37°C. Per mouse, 1-2 million cells were transferred (i.p.) and animals were injected with 67 ng pertussis toxin on day 0 (i.v.) and day 2 (i.p.) of EAE. Congenic animals (CD45.1 and CD45.2) were used to later distinguish endogenous and transferred T cells.

For Th17 adoptive transfer EAE, wildtype donor mice were actively immunized as described above and after 7 days their splenocytes lysed, cultured together with cells from dLNs for 3 days with IL-6 (5 ng/ml, Peprotech), hTGF-beta (0.25 ng/ml, Peprotech), anti-IFNg (14 ug/ml, Bio X Cell), IL-23 (6.5 ng/ml, Miltenyi) and MOG peptide (35 ug/ml) and sorted via positive selection of CD4^+^ T cells (MACS, Miltenyi). Per recipient, 3 million CD4^+^ T cells were transferred by iv. injection and animals were injected i.p. with pertussis toxin on day 0 and day 2 of EAE (300 ng).

Starting at the disease onset, approx. 8-10 days after induction of EAE, disease severity was monitored using an established scale from 0 to 4 to describe clinical symptoms: 0, no detectable signs of classical EAE; 0.5, paresis of distal limb tail; 1.0, tail paralysis; 1.5, hind-limb weakness; 2, hind-limb paresis; 2.5, partial hind-limb paralysis; 3, complete bilateral hind-limb paralysis; 3.5, ascending paralysis to torso; 4, paralysis of forelimbs and hind limbs. Mice with scores >3.5 were sacrificed. Dead animals were excluded or in warm exposure experiment given a score of 5 on death day due to increased deaths and high scores. Animals that reach a score >1.5 were provided with their chow diet, watered, inside the cage.

### B16-GMCSF tumor and thioglycolate elicited peritoneal inflammation

GMCSF-expressing B16 melanoma tumor cells (Dranoff et al., 1993) were cultured and s.c. engrafted as described (Menezes et al., 2016) into cold exposed and room temperature mice. Blood was drawn on several days after tumor implantation in heparin (Sigma) to analyze circulating monocyte expansion and phenotype using flow cytometry. For peritoneal inflammation, cold-exposed and room temperature mice were i.p. injected with 1ml of 4% thioglycolate (Sigma), as described before (Davies et al., 2013). Animals were sacrificed after 24h for analysis of peritoneal monocytes and monocyte derived cells using flow cytometry.

### FITC migration of DCs using skin painting of the flank

Mice were anesthetized with isoflourane, a small area of one flank (about 1×1cm) was shaved and 20 ul of FITC solution was applied, consisting of 1% FITC (Sigma-Aldrich: Fluorescein isothiocyanate isomer I) in Aceton/Dibutylphthalate (1:1, Sigma). After 12h and 18h, the 3 draining lymph nodes (axillary, brachial and inguinal) of ipsi- and contralateral side were harvested and cells harvested as described above.

### Antibody treatment for CCR2^+^ monocyte depletion

Monocytes were depleted by daily i.p. injections of anti-CCR2 mAb (clone: MC-21; 40 μg), starting one day before immunization on 8 consecutive days until sacrifice. Control mice received isotype rat IgG2b (Bio X Cell) at the same dose. Depletion efficiency was tested 2 days before EAE onset and sacrifice.

### Histology

For histologic analyses of spinal cord, mice were kept until 1-2 days after peak, perfused with PBS and subsequently 4% paraformaldehyde (PFA) and additionally immersion fixed in 4% PFA. Sectioning, embedding and staining were performed as previously described (Steinbach et al., 2016). Mice shown for histologic evaluation were control mice of an experiment and therefore injected with diphtheria toxin (Sigma). Stainings were developed with DAB (DAKO) and nuclei counterstained with hemalum (Merck). Sections were scanned using Pannoramic Digital Slide Scanner 250 FLASH II (3DHISTECH). Quantifications were performed blinded to the experimental condition. Anti-CD3 (polyclonal, DAKO) and anti-mac3 (M3/84, Biolegend) stainings were automatically quantified using Definiens Developer software with a custom-made script. Quantifications of Luxol Fast Blue (BDH laboratories)-Perdiodic Acid (Merck)-Schiff’s (Merck) (LFB-Pas) lesion areas, anti-APP (22C11, Millipore) and lesion number were performed manually using Pannoramic Viewer software (3DHISTECH). Number of lesions were counted in APP staining and only lesions were taken into account that were found in LFB-Pas as well. APP was quantified within lesions marked in LFB-Pas staining and reported per total white matter area. For representative LFB-Pas images, contrast was equally adjusted in the compared conditions in Adobe Photoshop.

For histologic analyses of adipose tissues, tissues were fixed in 4% paraformaldehyde (Sigma-Aldrich), paraffin embedded, cut in sections (5 μm) and stained with hematoxylin-eosin (H&E). Images were acquired using AxioScan Z1 (Zeiss) and the cell/lipid droplet size quantified with Qupath software (V0.2.2) using machine-learning classifiers.

### Immune cell isolation from tissues

Peripheral blood samples (10-30 ul) were obtained by facial vein puncture in 1 ul heparin (10U/ul, Sigma). Blood erythrocytes were lysed and cells fixed using BD FACS Lysing Solution. For other tissues that required red blood cell lysis, RBC Lysis buffer (Biolegend) was used. For the isolation of immune cells from the CNS, mice were anesthetized and transcardially perfused with PBS. Brain and spinal cord were minced, digested in DMEM (ThermoFisher) with Collagenase A (1mg/ml, Sigma) and DNaseI (0.1 mg/ml, Roche) for 1h at 37°C and homogenized using 70-μm cell strainers (BD). Leukocytes were separated using a discontinuous Percoll (GE healthcare) gradient (30% / 70%, 2500 RPM, 30 min, 4°C). Bone marrow cells were flushed from 1 femur per mouse. For analysis of DC and monocyte populations, axillary and brachial lymph nodes or spleen were cut into small pieces and digested in DMEM with collagenase D (1mg/ml, Roche) and DNaseI (10ug/ml, Roche) for 30-35 min at 37°C under constant pipetting. For studies of all other immune cells, lymph nodes or spleens were mashed through a 70 um cell strainer. For splenocytes, red blood cell lysis was performed (BioLegend).

### Flow cytometry

Dead cells were excluded from the analysis using Zombie NIR or Aqua Fixable Viability kit (BioLegend) for all tissues except blood. Surface staining was performed with directly labeled antibodies in FACS buffer (DPBS, 2.5% FCS, 10mM EDTA). Compensation controls were set up using Versacomp beads (Beckman Coulter). Antibodies were from BioLegend if not stated otherwise. For staining of bone marrow progenitors and mature monocytes, the following antibodies were used: Lin (CD3 (145-2C11), CD19 (6D5), NK1.1 (PK136), Ly6G (1A8), Ter-119), CD127 (A7R34), CD117 (ACK2, 2B8), CD115 (AFS98), CD135 (A2F10), Ly6C (HK1.4), CD11b (M1/70), Sca1 (D7), CD11c (N418), MHCII (M5/114.15.2). In figures, monocytes and monocyte-derived cells were abbreviated as “M” for simplicity reasons. Both Ly6C^hi^ monocytes and monocyte derived cells were defined mainly by their Ly6C expression (in addition to and exclusion of other markers), but irrespective of a co-expression of CD11c or pro/anti-inflammatory cytokines. “M” therefore comprises both, undifferentiated Ly6C^hi^ monocytes and monocytes that already gained APC or macrophage like functions but still express Ly6C. For characterization of blood and bone marrow monocytes, the following antibodies were used: CD45 (30-F11), Ly6G, CD11b, CD115, Ly6C, MHCII. For characterization of T cells in various tissues, the following antibodies were used: CD45, CD11b, Ly6G, CD4 (GK1.5), CD3, CD8 (53-6.7, BD), GM-CSF (MP1-22E9, ThermoFisher), IL-17 (TC11-18H10.1), IFNg (XMG1.2). Unspecific binding to FC receptors was blocked using anti-CD16/32 (BioLegend). Intracellular stainings were performed according to manufacturer’s instructions: Fixation/Intracellular Staining Permeabilization Wash Buffer (BioLegend) and Fixation buffer (BioLegend) were used for Arg1 (IC5868P, R&D), iNOS (CXNFT), IL-17, IFNg and GM-CSF. To assess for intracellular cytokine production of T cells, leukocytes were cultured for 4 h in the presence of PMA (end 50ng/ml, Sigma) and Ionomycin (end 1ug/ml) (Sigma), monensin (1:1000, BioLegend), and brefeldin A (1:1000, BioLegend). Absolute cell numbers were quantified using AccuCheck Counting Beads (Invitrogen). Flow cytometric samples were acquired on an Attune NxT (ThermoFisher) or LSRFortessa (BD) flow cytometer using compensation controls. FlowJo software (Treestar, Version 10) was used for analysis. Further antibodies used in DC analysis or high dimensional analysis, that were not yet mentioned before are CD49a (Ha31/8, BD), CD64 (X54-5/7.1), CX3CR1 (SA011F11), XCR1 (ZET), B220 (RA3-6B2), CD103 (M290, BD), F4/80 (BM8), CD49d (R1-2), PDCA-1 (927), Siglec-F (E50-2440, BD). High-dimensional flow cytometry analysis was performed using BD FACSymphony. Raw data was pre-processed using FlowJo followed by transformation, percentile normalization in R, dimensionality reduction and visualization by Uniform Manifold Approximation and Projection (UMAP) (McInnes et al., 2018). The FlowSOM algorithm was used for automated clustering (Van Gassen et al., 2015) using UMAP with overlaid mean marker expression values and a heatmap of median expression values (Hartmann et al., 2016; Mrdjen et al., 2018). Data shown in Figure 2C is the same as in Figure 5E but was visualized together with data from other time points to highlight shifts of populations over the time in Figure 5E.

#### Oxygen Consumption Rate (OCR)

OCR was determined as before (Dunham-Snary et al., 2014), in 3.2-8mg punches (2.5 mm) of adipose tissue. After washing in wash buffer (25mM HEPES, 1% fatty acid-free BSA, 2.5 mM glucose, in salt solution: 4mM KCl, 115 mM NaCl, 1 mM MgCl2, 1.5 mM KH2PO4, 2 mM MgSO4, 1.5 mM CaCl2 at pH 7.4), tissue punches were placed in islet capture plates (model XF24, Seahorse Bioscience) and washed with wash and subsequently with respiration buffer (0.1% fatty acid-free BSA, 10 mM glucose, 2 mM L-glutamine, 0.5 mM L-carnitine, 1 mM sodium pyruvate in salt solution at pH 7.4), in which they were further equilibrated for 50 or 60 min at 37°C. Samples were analyzed using a Seahorse XF24 Analyzer (Seahorse Bioscience) and Wave software with 3 minutes mix; 2 minutes rest; 3 minutes measurements in basal condition.

### RNA Extraction for RNA sequencing and qPCR

After preparation from mice, spinal cord and adipose tissues were stored at −80°C until further processed. For RNA isolation, frozen tissues were mechanically homogenized in 1 ml Trizol (Thermo Fisher Scientific) with 1 stainless steel bead (5 mm) per sample using the TissueLyserII (Qiagen) by shaking for 50 s at 30 Hz. 200 μL chloroform was added to homogenized samples in trizol, followed by shaking for 15 s and centrifugation (15 min, 12000 RCF, 4°C). The chloroform phase was collected, shaken for 15 s with 500 μL isopropanol and centrifuged (15 min, 12000 RCF, 4°C). The pellet obtained was washed twice with 70% ethanol (10 min, 8000 RCF, 4°C) and resuspended in 50 μL PCR-grade water. Bone marrows were flushed right after sectioning from mouse, cells centrifuged, loaded onto shredder columns (Qiagen). Shredded bone marrow cells or FACS sorted cells were frozen (−80°C) in RLT buffer with 1% beta-mercaptoethanol until RNA extraction via RNA mini kit or micro kit (Qiagen), respectively. For Sequencing, RNA integrity number (RIN) was determined (Bioanalyzer 2100, Agilent). For qPCR, RNA concentration and purity were determined (NanoDrop, Thermo Fisher Scientific) and 1 μg RNA was used per sample for retrotranscription using the High-Capacity cDNA Reverse Transcription Kit (Applied Biosystems). Expression of mRNA was measured by qPCR on a LightCycler 480 II (Roche) with PowerUp SYBR Green Master Mix (Applied Biosystems). Relative mRNA expression was normalized to at least two house keeping genes and to room temperature control mice.

### Blood monocyte purification for RNA sequencing

Blood was taken retro-orbital under terminal anesthesia in polypropylen facs tubes containing 7ul heparin (10U/ul) and coated with 100 IU/mL heparin. Anti-CD115-PE was directly added and MACS separation was performed using Anti-PE MicroBeads (Miltenyi) and AutoMACS Pro Seperator (Miltenyi). Immediately after, monocytes were sorted as CD115+, DAPI-(ThermoFisher) cells (FACSAriaII, BD, Bio-Rad S3 or MoFlo Astrios, Beckman Coulter).

### RNAseq sequencing and analysis

mRNA sequencing was performed at the iGE3 Genomics Platform at the CMU of the University of Geneva. Libraries for sequencing were prepared with the TruSeq stranded mRNA kit (Illumina) for bone marrow and spinal cord. The SMART-Seq v4 ultra-low input kit (Clontech) combined with the Nextera kit (Illumina) were used respectively for cDNA amplification and library preparation for monocytes (steady state and EAE). Bone marrow, spinal cord, and monocytes (EAE) were sequenced with read length SR50 and steady state monocytes with read length SR100 (Illumina HiSeq 4000). The sequencing quality control was done with FastQC v.0.11.5 (http://www.bioinformatics.babraham.ac.uk/projects/fastqc/). The reads were mapped with STAR aligner (Dobin et al., 2013) to the UCSC Mus musculus mm10 reference. The transcriptome metrics were evaluated with the Picard tools v.1.141 (http://picard.sourceforge.net/). The table of counts with the number of reads mapping to each gene feature of the Mus musculus mm10 reference was prepared with HTSeq v0.9.1 (htseq-count, http://www-huber.embl.de/users/anders/HTSeq/). Raw counts were then further processed and analyzed by R/Bioconductor package EdgeR v. 3.4.2 (McCarthy et al., 2012a), for differential expression analysis. The counts were normalized according to the library size and filtered. Only genes having log count per million reads (cpm)>0 were kept for further analyses. After normalization of the counts, transcript abundances were compared in pairwise conditions in a modified Fischer exact test (as implemented in edgeR). The differentially expressed genes p-values were corrected for multiple testing error with a 5% FDR (false discovery rate). The correction used is Benjamini-Hochberg (BH). Transcripts with log(cpm)>0 and p≤0.05 were subsequently subjected to pathway analysis. For the relativeness and reactome analysis, genes were called significantly differentially expressed between any given two conditions when p<0.05 and FC>2. Reactome pathways database was used with ReactomePA R-package (Yu and He, 2016), reporting the enrichment ratio (# DEGs/Total Genes in the dataset) and FDR-adjusted Pvalue computed by Fisher exact test. For gene ontology analysis, R/Bioconductor package topGO (https://bioconductor.org/packages/release/bioc/vignettes/topGO/inst/doc/topGO.pdf), together with Rgraphviz was used. MetaCore (Clarivate Analytics) Pathway Maps, Gene Ontology and Metabolic Networks were used.

### Statistical analysis

To assess significant differences, GraphPad 8.0 (Prism) was used. For two groups, two-tailed Student’s t test was used. For more than two groups, multiple t-tests were applied with Holm-Sidak correction. Two-waxy ANOVA was used to compare two variables between groups. Mantel-Cox was applied to test the incidence or survival distributions of two groups. P < 0.05 was considered significant (*), p < 0.01 (**), p < 0.001 (***).

